# Female gametophyte expressed *Arabidopsis thaliana* lipid transfer proteins AtLtpI.4 and AtLtpI.8 provide a link between callose homeostasis, pollen tube guidance, and fertilization success

**DOI:** 10.1101/2021.01.13.426551

**Authors:** Khushbu Kumari, Meng Zhao, Sebastian Britz, Christine Weiste, Wolfgang Dröge-Laser, Christian Stigloher, Rosalia Deeken, Dirk Becker

## Abstract

Non-specific lipid transfer proteins (LTPs) represent a sub-class among the large family of Cysteine-rich proteins (CRPs) specific to land plants. LTPs possess a hydrophobic cavity, enabling them to bind and stabilize a variety of lipid molecules outside membranes. In line with the existence of an N-terminal signal peptide, secreted LTPs represent a well-suited mobile signal carrier in the plant’s extracellular matrix. Thus, LTPs are currently considered as key players to mediate the bulk flow of lipids between membranes/compartments as well as the buildup of lipid barrier polymers including cutin and suberin.

Here, we show that floral expressed *Arabidopsis thaliana* AtLtpI.4 (AtLTP2) and AtLtpI.8 (AtLTP5), mutually control cell-cell communication between growing pollen tubes and ovules during fertilization. Arabidopsis mutants lacking functional AtLtpI.4 and AtLtpI.8 exhibit significantly reduced fertilization success. Cross-pollination and cell biological analyses revealed that *AtLtpI.4/I.8* double mutants are impaired in pollen tube guidance towards ovules. Our finding that the *AtLtpI.4/I.8* phenotype correlates with aberrant callose depositions in the micropylar region during ovule development suggests that both LTPs represent novel players of a joint signaling pathway that controls callose homeostasis in the female gametophyte.

## INTRODUCTION

The evolution of apoplastic diffusion barriers in land plants is represented as one of the key molecular adaptations during their transition to land (Nawrath et al., 2013). The diverse physiological functions of these apoplastic interfaces depend on their composition, structure and properties. Within the cellulose matrix of the cell wall polysaccharides such as pectins or callose can serve as apoplastic barriers, restricting the diffusion of high molecular weight molecules. The ß-glucan callose is composed of glucose residues linked together through β-1,3-linkages and it acts as a dynamic, temporary barrier. Membrane localized callose synthases (CalSs) synthesize, while secreted, extracellular β-1,3-glucanases (BGs) degrade callose directly on the spot. In response to pathogen attack callose deposition provides for rapid closure of plasmodesmata (Cheval et al., 2020). In addition, callose is known to be involved in pollen– ovule interaction during fertilization (De Martinis and Mariani, 1999). It has been shown that abnormal callose formation at the micropylar region prevented normal pollen tube guidance in *Arabidopsis thaliana* flowers (Kocsis and Jakab, 2008). The signaling pathways regulating callose deposition or its removal are, however, largely unknown.

Polymers based on lipid polyesters or organic compounds such as cutin or suberin represent hydrophobic cell wall reinforcements (Philippe et al., 2020). From a developmental perspective for instance they prevent organ fusion and thus control plant morphology (Pollard et al., 2008). Secreted by epidermal cells, the hydrophobic, cutin containing plant cuticle, covers the shoot surface. Thereby, it prevents land plants from excess water loss and at the same time provides for a protective layer against abiotic and biotic stresses. On the other hand, suberin layers are mostly deposited on the inner layer of cell walls. It is found, for example, in tubers and seed coats as well as in root endodermal cells where suberin acts as a diffusion barrier for selective nutrient uptake by enforcing symplastic nutrient transport (Baxter et al., 2009).

Cutin is mainly composed of derivatives of C16 and C18 fatty acids while suberin is largely derived from long-chain aliphatic acids (>C16), fatty alcohols, hydroxycinnamic acids and glycerol. C16:0, C18:0 and C18:1 fatty acids represent the initial lipid precursors of both cutin and suberin. While ATP binding cassette (ABC) transporters of the ABCG subfamily seem to mediate the transport of suberin monomers across the plasma membrane, nonspecific lipid transfer proteins (LTPs) have been suggested to provide for the movement of lipid polymer and wax components to the sites of polymer accumulation on the extracellular side of the plasma membrane (Edqvist et al., 2018).

LTPs are small Cysteine-rich proteins (CRPs). They usually possess an N-terminal signal peptide as well as an even number of at least four or six cysteine residues, while a specific cysteine pattern is characteristic for a given LTP subfamily (Silverstein et al., 2007; Huang et al., 2015). Their physiological functions include membrane biogenesis, cell differentiation, intercellular and extracellular signaling, as well as lipid barrier formation (Boutrot et al., 2008; Cecchini et al., 2015; Deeken et al., 2016). In LTPs, an eight-Cys motif (8CM; with the general sequence C-Xn-C-Xn-CC-Xn-CXC-Xn-C-Xn-C) results in the formation of four conserved disulfide bridges. Thereby, LTPs build a central hydrophobic cleft for the binding of hydrophobic ligands, including fatty acids and other lipids. Based on transcriptomic profiling as well as gene-regulatory network analyses, it appears that LTPs in angiosperms can be classified into root, shoot, and reproductive LTPs (Salminen et al., 2016). This finding is in line with recent studies, demonstrating that of the 756 CRP-encoding genes present in the *Arabidopsis thaliana* genome, 205 and 139 CRP genes were reported as specifically expressed in the female and male gametophyte, respectively with many of them up-regulated upon pollination (Huang et al., 2015; Mondragón-Palomino et al., 2017). It further suggests, that extracellular Cysteine-rich proteins (CRPs) together with their corresponding membrane receptors seem to act as key players in male and female cell-to-cell communication in angiosperms (Bircheneder and Dresselhaus, 2016). Secreted by the synergid cells, LURE proteins, belonging to the Defensin-like (DEFLs) class of CRPs, were the first molecules identified in *T. fournieri* (Okuda et al., 2009) and later also in *A. thaliana* and *A. lyrata* (Takeuchi and Higashiyama, 2012) to mediate the attraction of pollen tubes towards ovules. In *Arabidopsis thaliana*, LURE1 recognition is mediated by multiple pollen expressed receptorlike kinases (RLKs) that seem to act as functionally redundant (Zhang et al., 2017). Notably, in maize, the small egg apparatus expressed, non-CRP peptide EGG APPARATUS1 (EA1) serves as a pollen tube attractant (Uebler et al., 2015), while the DEFLs EMBRYO SAC1-4 (ES1-4) trigger pollen tube burst, but do not possess guidance activity (Amien et al., 2010). In *A. thaliana*, however, pollen tube burst is controlled by cysteine-rich peptides of the Rapid Alkalinization Factor (RALF) family. They serve as ligands for their respective receptors of the *Catharanthus roseus* Receptor-Like Kinase 1-like (CrRLK1L) family and the competition of male and female-derived RALF peptides provides for pollen tube integrity during growth and sperm cell release, respectively (Ge et al., 2017).

In comparison to LURE or RALF peptides, little is known about the role of ‘reproductive’ LTPs in the fertilization processes. In *Lilium longiflorum*, the LlLTP1.1 (SCA) is secreted from the pistil transmitting tract epidermis and was shown to regulate the formation of an adhesive matrix together with pectin that guides the pollen tubes to the ovules (Park and Lord, 2003). Of the seven SCA-like LTPs in *Arabidopsis thaliana*, a gain-of-function mutant of AtLtp1.8 (LTP5; *ltp5-1*) exhibited defects in pollination and seed formation, characterized by delayed pollen tube growth and a decreased number of fertilized ovules. The corresponding AtLTP1.8 loss-of-function mutant, however, appeared wild-type-like (Chae et al., 2009; Chae et al., 2010), suggesting that Arabidopsis AtLTP1.8 acts in concert with other ‘reproductive’ LTPs. The existence of an apoplastic lipid barrier between the maternal inner integument and the nucellus in developing *Arabidopsis thaliana* seeds though indicates that ovule development and fertilization is based on an intricate communication between cells of the female and male gametophyte as well as maternal tissues (Loubery et al., 2018; Coen et al., 2019). To answer the question, whether ‘reproductive’ LTPs in *Arabidopsis thaliana* have an impact on the formation of these interfaces, we identified type-I LTPs co-expressed in the *Arabidopsis thaliana* female gametophyte and analyzed their putative role in fertilization.

## RESULTS

### Expression patterns of Type-I Arabidopsis LTP genes in reproductive and vegetative tissues

To get a first overview about LTP expression in the Arabidopsis flower with a special emphasis on the female gametophyte we used publicly available expression data bases including TAIR (www.arabidopsis.org), genevestigator (genevestigator.com/gv/) and ePlant (bar.utoronto.ca/eplant/). We identified AtLTP1.5 (**AtLTP1**; AT2G3854), AtLTP1.4 (**AtLTP2**; AT2G38530), AtLTP1.12 (**AtLTP3**; AT5G59320), AtLTP1.11 (**AtLTP4**; AT5G59310), AtLTP1.8 (**AtLTP5**; AT3G51600), AtLTP1.6 (**AtLTP6**; AT3G08770), and AtLTP1.7 (**AtLTP12**; AT3G51590) to be highly expressed in the stigma, pistil and/or ovule (see **Supplemental Table 1**).

To be able to follow the spatial and temporal expression of these LTPs more precisely, we generated stable *A. thaliana* plants expressing transcriptional fusions of individual LTP-promoters (~2000 bp; see methods) and the beta-glucuronidase (GUS) reporter coding sequence. On the whole seedling level all tested promoters were active in leaves and roots (**Supplemental Figure 1**). In floral and reproductive tissues GUS expression driven by the *AtLTP1.4* promoter was detected in style, stigma, funiculus and transmitting tract (**Figure 1A and 1B**). The *AtLTP1.12* (**Figure 1C**), *AtLTP1.11* (**Figure 1D**), *AtLTP1.6* (**Figure 1G**) *and AtLTP1.7* (**Figure 1H**) genes were also expressed in the style, in anther filaments and sepals, while *AtLTP1.8* was expressed in almost all floral tissues (**Figure 1E and 1F**), including the stigmas of young buds and mature flower, the style, anther, anther filament, ovule, the transmitting tract and the funiculus. Consistently, the selected *Arabidopsis thaliana* LTPs showed high level of expression in the pistil, specifically in the style.

**Figure 1.**
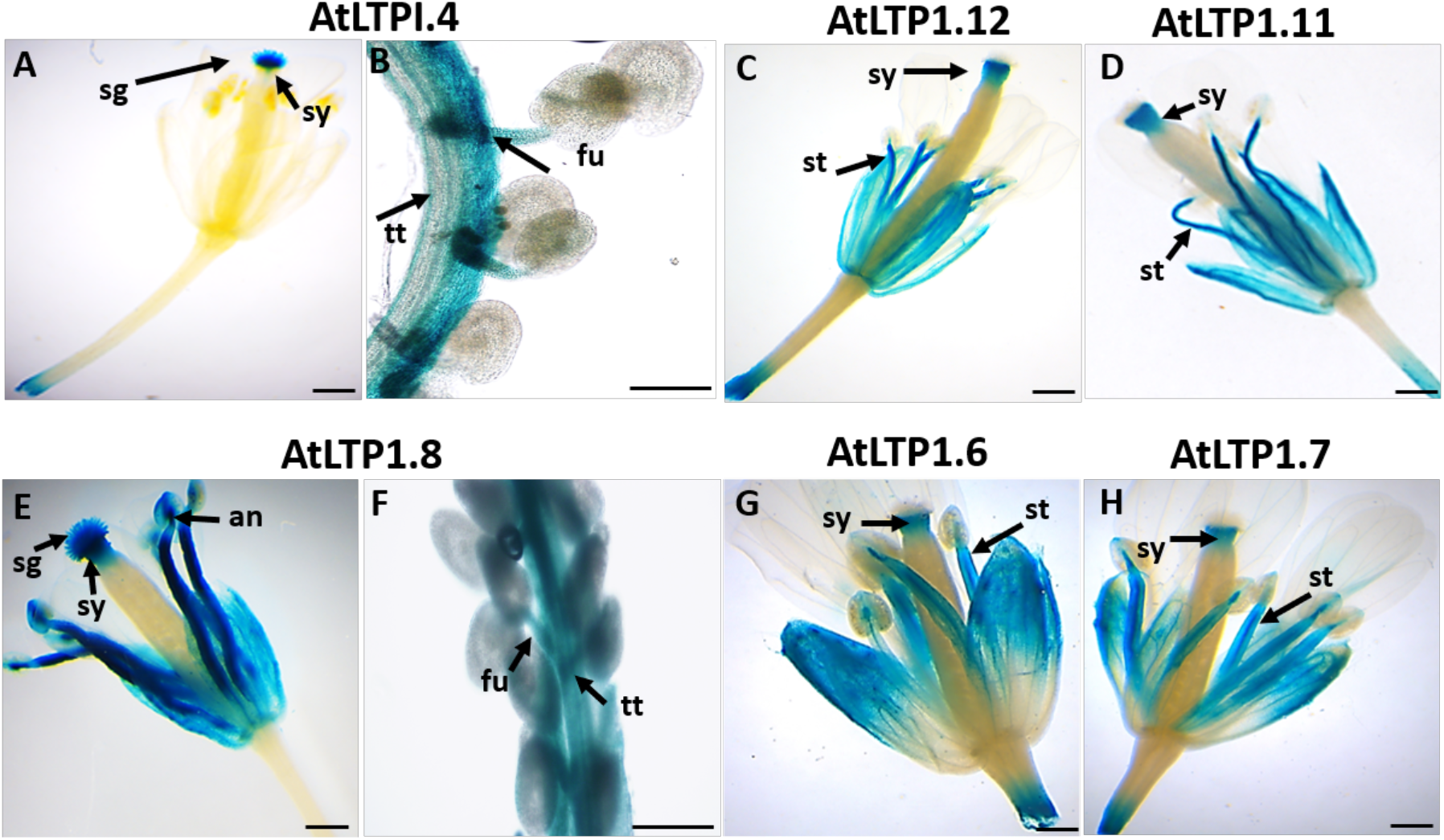
GUS Expression of 6 LTPs in the floral and reproductive tissue of *At*. **(A)** *AtLTP1.4* gene expression in the stigma and style, **(B)** funiculus and transmitting tract. **(C)** *AtLTP1.12* gene expression in the style and anther filaments. **(D)** *AtLTP1.11* gene expression in the style and anther filaments. **(E)** *AtLTP1.8* gene expression in the stigma, style, anther and anther filament, **(F)** funiculus and transmitting tract. **(G)** *AtLTP1.6* gene expression in the style and anther filaments. **(H)** *AtLTP1.7* gene expression in the style and anther filaments. Plants were grown under long-day condition (16hrs light/ 8hrs dark). GUS signals were developed for 2:30hrs. **sg**-sigma, **sy**-style, **st**-stamen, **fu**-funiculus, **tt**-transmitting tract, **an**-anthers. A, C, D, E, F, G, H; Scale: 2mm, B, F; Scale: 50μm.

### Distinct subcellular localization patterns characterize individual AtLTPs

The presence of a predicted N-terminal signal peptide (21–27 amino acids in length) suggests, that all seven *Arabidopsis thaliana* LTPs under investigation are most likely entering the cellular secretory pathways. To disclose their subcellular localization we transiently expressed individual AtLTPs fused to the Yellow Fluorescent Protein mVenus together with the cell-wall marker AtLTP1.5::mCherry (Deeken et al., 2016) in the *N. benthamiana* system. Co-expression of either AtLTP1.4::YFP (**Figure 2A** upper row), AtLTP1.8::YFP (**Figure 2B** upper row) or AtLTP1.6::YFP (**Supplemental Figure 2A**) with AtLTP1.5::mCherry clearly revealed their co-localization at the cell periphery. Upon plasmolysis, both, YFP and mCherry fluorescence co-localized in the apoplastic space (**Figure 2A, 2B** and **Supplemental Figure 2A** middle and lower row). This observation was supported by the Pearson correlation coefficients ≥ 0.9 for the co-localization of AtLTP1.4, AtLTP1.8 and AtLTP1.6 with AtLTP1.5 (**Supplemental Figure 2E**) and further by *Correlative Light and Electron Microscopy* (CLEM) as exemplified for AtLTP1.8. Following an immunostaining of ultrathin sections from AtLTP1.8::YFP expressing leaves using an anti-GFP antibody, *Structured Illumination Microscopy* (SIM) (**Figure 3A, 3D, 3G**) and subsequent *Scanning Electron Microscopy* (SEM) imaging (**Figure 3B, 3E, 3H**) localized the AtLTP1.8::YFP signal in the extracellular space after correlation (**Figure 3C, 3F, 3I)**.

**Figure 2.**
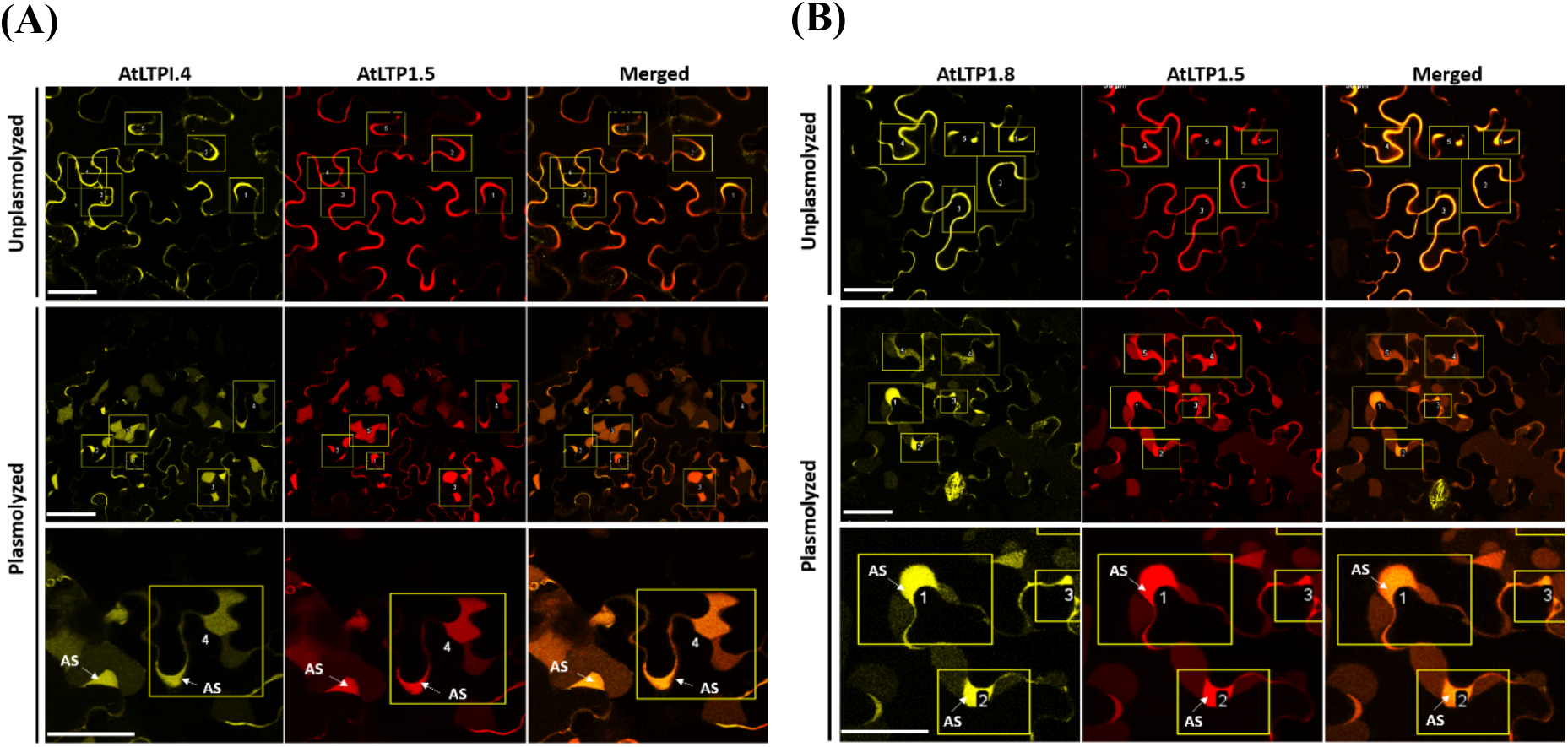
Sub-cellular localization of AtLTP1.4 and AtLTP1.8. **(A)** AtLTPI.4 **(B)** AtLTP1.8 fused to YFP in *N. benthamiana* epidermal cells imaged four days post infiltration. Coexpression of the two-fusion proteins 35SLTP1.4::YFP/ 35SLTP1.8::YFP and cell wall marker 35SLTP1.5::mCherry (upper row); Scale: 50μm. Co-infiltrated leaves were plasmolyzed with 0.5M KNO_3_and documented after 5mins of incubation (middle row); Scale: 50μm. Higher magnified images depicting co-localization of 35SLTP1.4::YFP/35SLTP1.8::YFP with 35SLTP1.5::mCherry in the apoplastic space (AS) of the plasmolyzed cell (lower row); Scale:50μm. Each selected region of interest (ROI) is marked in yellow box and numbered from 1-5. mCherry (Excitation: 561 nm, Emission: 580–615 nm); mVenus (Excitation: 514 nm, Emission: 530–555 nm).

**Figure 3.**
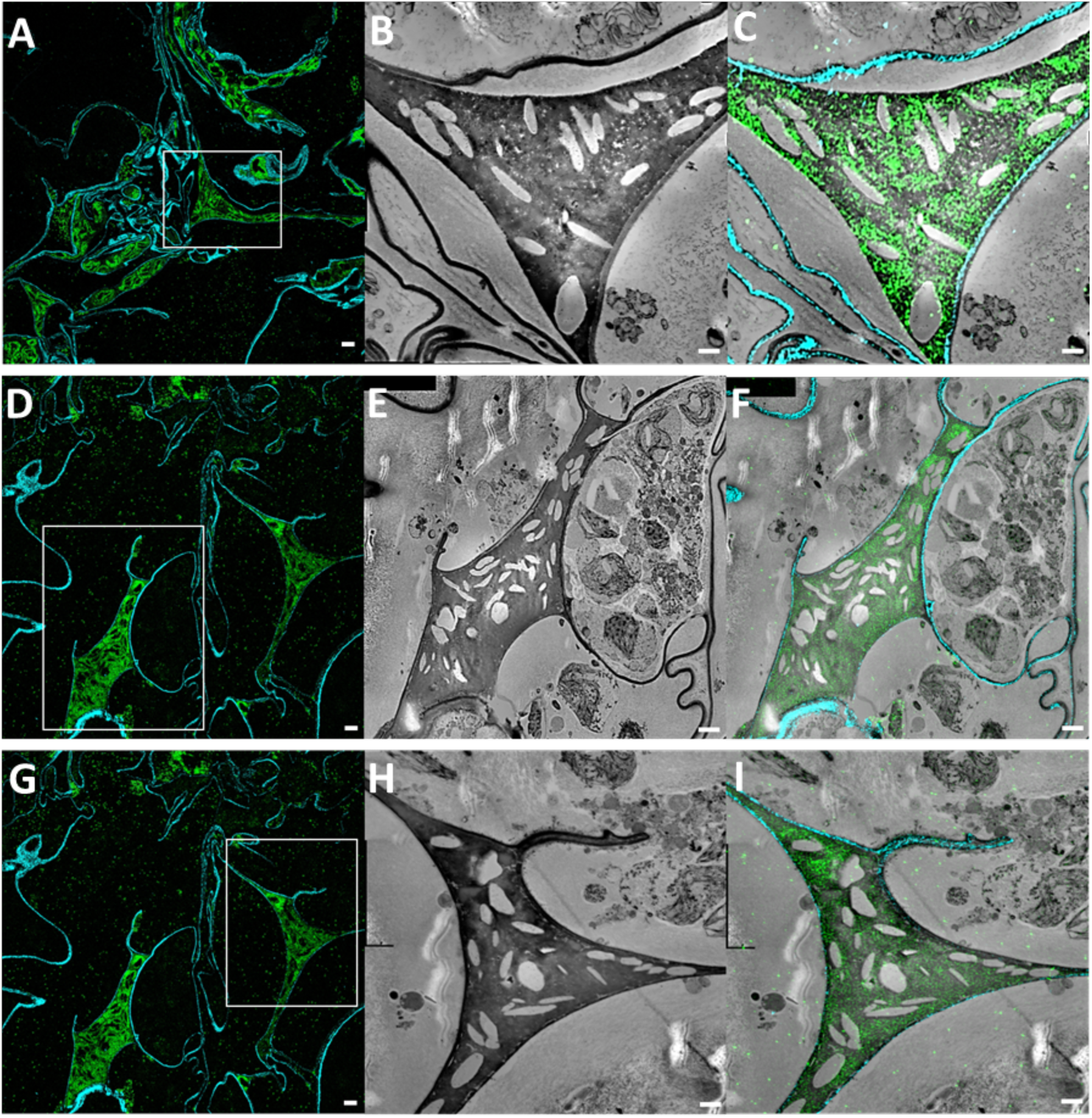
Localization of AtLTP1.8::YFP using super-resolution CLEM on ultrathin sections of *N. benthamiana* agro-infiltrated leaf samples. 100 nm LR-White section of an infiltrated tobacco leaf sample imaged with a *Structured Illumination Microscopy* (SIM) (A, D, G) and with the *Scanning Electron Microscope* (SEM) (B, E, H). Same images as in (B, E, H) overlaid with SIM fluorescence channels; Scale:1μm (C, F, I). Cell wall (turquoise blue) and extracellular space (green); Scale:1μm.

In contrast, AtLTP1.7, AtLTP1.11 and AtLTP1.12 did not exhibit co-localization with the cellwall marker AtLTP1.5::mCherry (**Supplemental Figure 2**). Upon plasmolysis, AtLTP1.7::YFP, AtLTP1.11::YFP and AtLTP1.12::YFP fluorescence was progressively detached from the cell-wall and was associated with the shrinking plasma membrane and hechtian strands. At the same time, co-expressed AtLTP1.5::mCherry was clearly detectable in the apoplastic compartment delimited by the plasma membrane (**Supplemental Figure 2B, 2C, 2D**). In line with this observation, the Pearson correlation coefficient for co-localization of AtLTP1.7, AtLTP1.11 and AtLTP1.12 with AtLTP1.5 was ≤ 0.4 (**Supplemental Figure 2E**). Taken together, these results indicate that AtLTP1.7, AtLTP1.11 and AtLTP1.12 are associated with the plasma membrane, while AtLTP1.4, AtLTP1.8 and AtLTP1.6 reside in the apoplast. Next, we performed quantitative PCR (qPCR) and monitored LTP expression in pistils in response to pollination. With the exception of AtLTP1.7, the female gametophyte expressing AtLTPs were significantly induced upon pollination, corroborating their putative role during fertilization (**Figure 4**).

**Figure 4.**
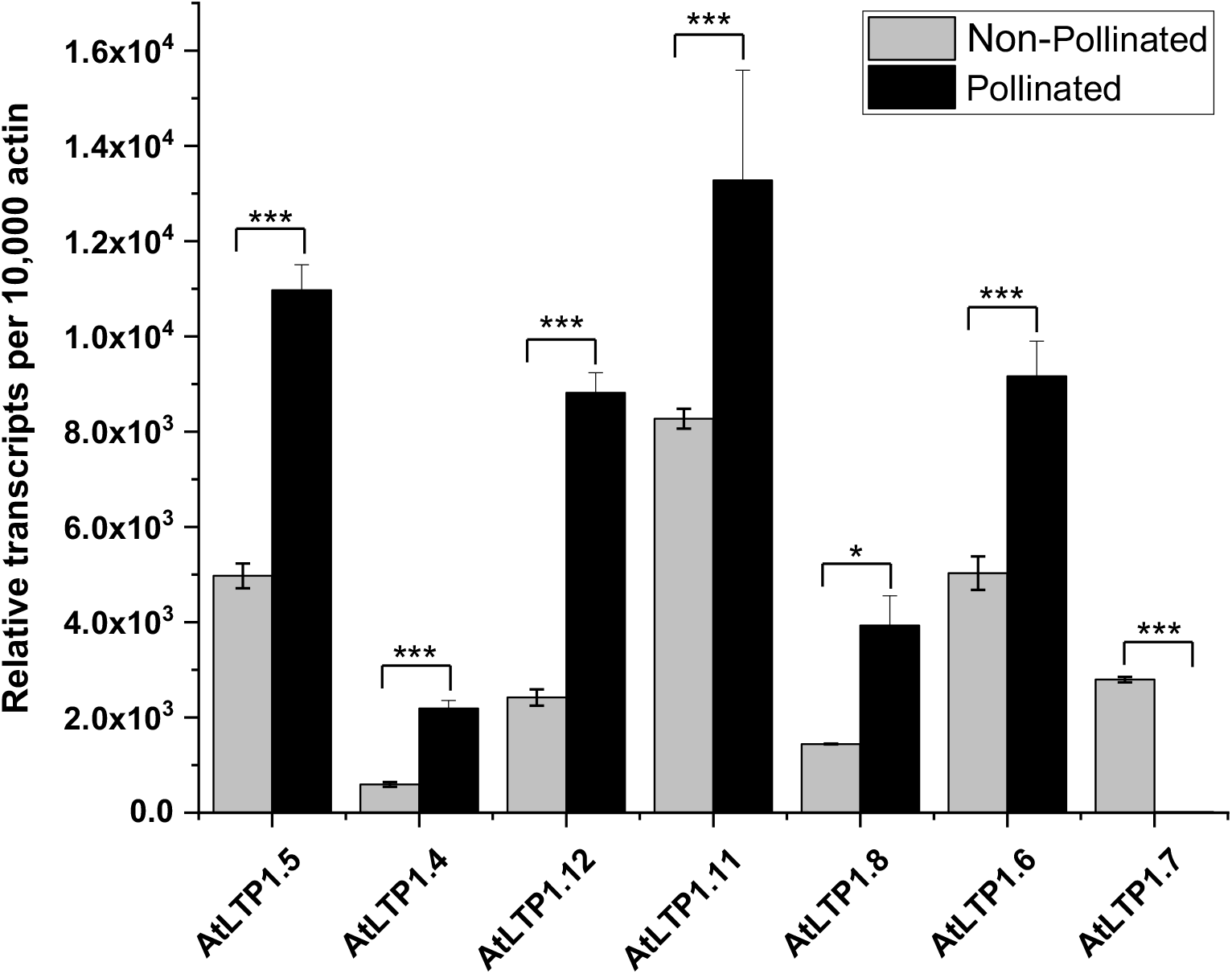
Transcriptional regulation of LTPs upon pollination. LTPs (*AtLTP1.5 (AtLTP1), AtLTPI.4 (AtLTP2), AtLTP1.12 (AtLTP3), AtLTP1.11 (AtLTP4), AtLTP1.8 (AtLTP5), AtLTP1.6 (AtLTP6), and AtLTP1.7 (AtLTP12)*) gene expression relative to actin gene analyzed by qRT-PCR comparing pollinated and non-pollinated pistils of Arabidopsis (WT, Col-0). Three independent biological replicates were used for each treatment. Data are shown as mean ± SD. Asterisks indicate significance differences in pollinated pistils compared to non-pollinated pistils, n=3; *, *P*<0.05, **, *P*<0.01, ***, *P*<0.001 (*t*-test).

### The role of LTPs in fertilization

Following this assumption, we identified and characterized T-DNA insertion mutants for flower expressed *Arabidopsis thaliana* LTPs. All mutant lines were confirmed to be homozygous for T-DNA insertion (**Supplemental Figure 3A**). Gene expression analysis on flower tissue however revealed, that only AtLTP1.4 (GABI_639E08), AtLTP1.12 (SALK_095428C), AtLTP1.8 (FLAG_529A03; SALK_104674), AtLTP1.6 (SALK_120555) and AtLTP1.7 (FLAG_487D10) were true knock-outs, since their transcripts were not detectable by qPCR (**Supplemental Figure 3B**). Although less expressed, when compared to wild type, transcripts for AtLTP1.11 (GABI_194C08, GABI_172E03) and AtLTP1.8 (SALK_104674) were clearly detectable, suggesting that these mutant lines represent knock-downs. With the exception of the previously published gain-of-function AtLTP1.8 mutant (SALK_104674) (Chae et al., 2009; Chae et al., 2010), no discernible phenotype was evident for any of the other single homozygous mutant lines on the basis of pollen germination, *in vivo* pollen tube growth and seed set **(Supplemental Figure 3C)**. This might be attributable to functional redundancy of highly homologous LTPs.

Since phylogenetic analyses revealed that AtLTP1.4 and AtLTP1.8 are highly homologous and belong to the same branch within the type-I LTP subclade (**Supplemental Figure 4A**) (Boutrot et al., 2008) we therefore decided to generate a *ltp1.4/1.8* double knock-out mutant by employing the recently published germ-line egg cell-specific promoter-controlled EPC CRISPR/Cas9 system (Wang et al., 2015) allowing targeted mutation of two genes simultaneously. Guide RNAs were designed to target the first exon of LTP1.4 and LTP1.8 (**Supplemental Figure 5A**). Following PCR-based genotyping and Sanger sequencing, four transgenic lines (#P9-P2-P2, #P9-P3-P3, #*ltp1.4/1.8-1* and #*ltp1.4/1.8-2*) were selected for further high throughput sequencing analysis (**Supplemental Figure 5B** and **5C**). Sequencing analysis of more than 2,000,000 reads using the CRISPresso computational tool (Pinello et al., 2016; Canver et al., 2018) revealed that #*ltp1.4/1.8-1* and #*ltp1.4/1.8-2* exhibits the highest mutation rates displaying 99.8% and 99.9% frameshift mutations in AtLTP1.4 and AtLTP1.8 respectively (**Supplemental Figure 5D**). Based on indel mutations in both, AtLTP1.4 and AtLTP1.8, the resulting frameshifts, either gave rise to premature stop codons or altered proteins sequences right next to the guide RNA (**Supplemental Figure 5C**). In line with these frameshifts subsequent qPCR analysis in pistils revealed that AtLTP1.4 as well as AtLTP1.8 expression was absent in both lines (**Supplemental Figure 5E and 5F**).

### *The ltp1.4/1.8* double knock-out mutant exhibits fertilization defects and impaired pollen tube guidance

Upon phenotypic characterization of the two *ltp1.4/1.8* double mutant lines, we observed, that plants lacking functional AtLTP1.4 and AtLTP1.8 proteins exhibited severe fertilization defects (**Figure 5**). When compared to wild-type, siliques of both *ltp1.4/1.8* double mutant lines were significantly shorter (**Figure 5A** and **5B**). In addition, mutant lines exhibited an increased number of non-fertilized, aborted ovules (**Figure 5A** and **5C**), which resulted in reduced seed formation (**Figure 5A** and **5D**). In contrast to the established AtLTP1.8 gain-of-function mutant phenotype, in *ltp1.4/1.8* double mutants, unfertilized ovules were randomly distributed along the entire ovary and were not restricted to its base (Chae et al., 2009). To elucidate, whether this phenotype in *ltp1.4/1.8* double mutants is caused by impaired pollen tube growth or compromised pollen-pistil interaction, we followed *in vivo* pollen tube growth in pistils of self-pollinated mature flowers of stage 12 of the *ltp1.4/1.8* double mutant lines. According to aniline-blue staining and in contrast to the AtLTP1.8 gain-of-function mutant, *ltp1.4/1.8* double knockout pollen was competent in germination and displayed wild-type-like tube growth along the entire transmitting tract *in vivo* (**Figure 5E**). In wild-type ovaries, the micropylar region of ovules were successfully targeted by single pollen tubes with high efficiency (**Figure 5E, left**). In *ltp1.4/1.8* mutant lines, however, pollen tubes failed to emerge from the transmitting tract and did not turn towards the micropyle (**Figure 5E, middle** and **right)**, suggesting that the apparent pollen tube guidance phenotype might be attributable to compromised functions of the female rather than the male gametophyte.

**Figure 5.**
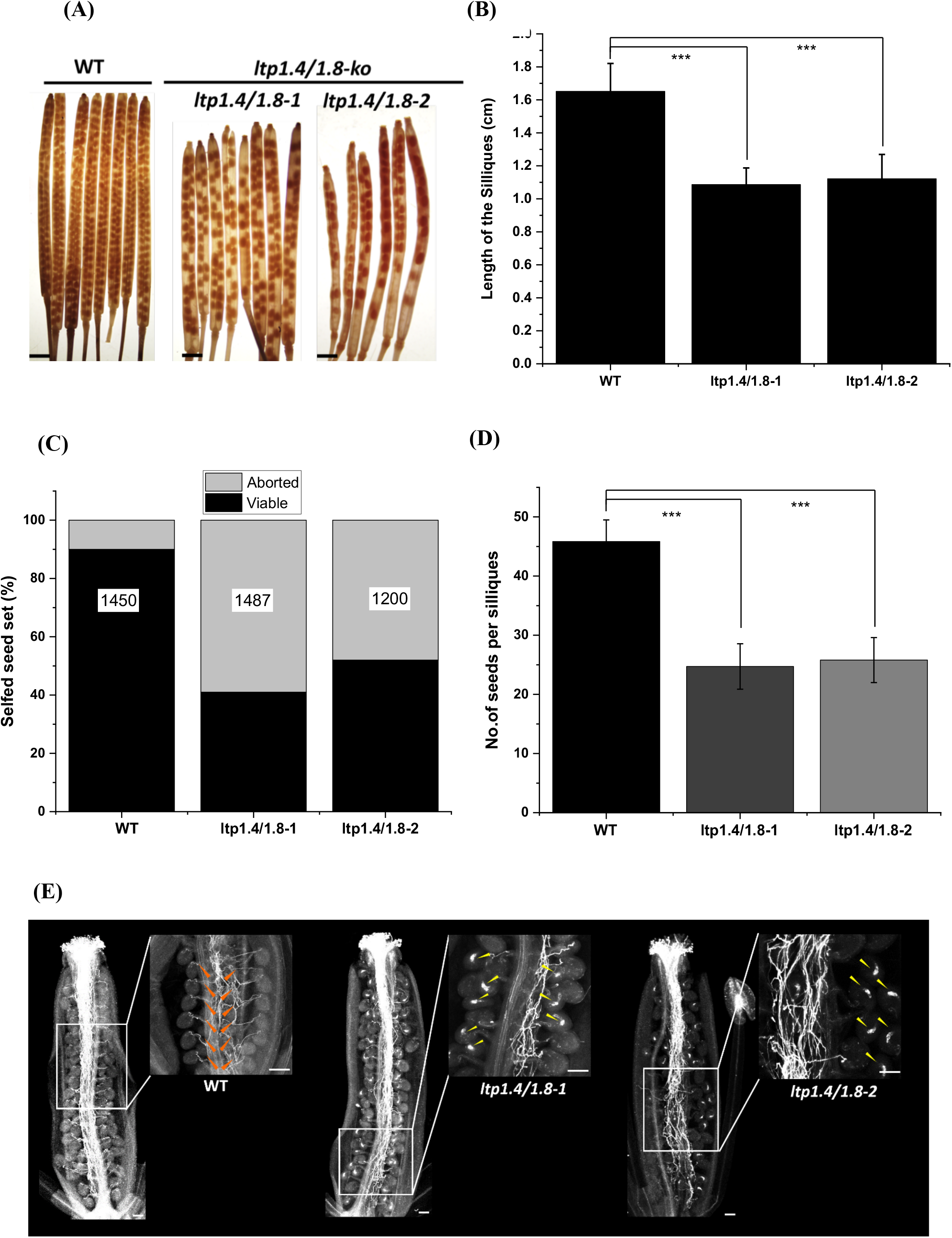
Characterization of *ltp1.4/1.8-1* double knock-out mutant. The *ltp1.4/1.8* mutant were semi-sterile and showed reduce seed set due to defect in the female gametophyte**. (A)** Siliques of the wild type are filled with seeds, whereas *ltp1.4/1.8-1* and *ltp1.4/1.8-2* mutant siliques showed severe seed abortion. Mature siliques were de-stained with 100% ethanol to examine seed sets (n = 60); Scale bars = 1mm. **(B)** Sizes of mature siliques from self-pollinated plants were analyzed. **(C-D)** The reduced seed number in *ltp1.4/1.8-1* and *ltp1.4/1.8-2* mutants in comparison to the wild-type. Total number of seeds analyzed are in the center of each column. **(E)** Defective pollen tube guidance and decreased ovule-targeting ability observed in the *ltp1.4/1.8-1* and *ltp1.4/1.8-2* mutant lines. Confocal fluorescence microscopy images of self-pollinated wild type and *l ltp1.4/1.8-1* and *ltp1.4/1.8-2* mutant pistils emasculated from stage 12 mature flower. **(E-left)** The pollen tubes were attracted normally to the wild-type ovules in the self-pollinated wild-type pistil (indicated by orange arrowhead), whereas in selfpollinated mutant **(E-middle)** *ltp1.4/1.8-1* and **(E-right)** *ltp1.4/1.8-2* the callosized ovules failed to attract the pollen tubes toward its micropyle (indicated by yellow arrowhead). Data are shown as mean ± SD. The error bars and asterisks indicate values that differ significantly from those of the wild-type plant; *, *P*< 0.05; **, *P*< 0.01; and ***, *P*< 0.001 (*t* test).

To answer this question, we therefore performed reciprocal cross pollination experiments between wild-type and *ltp1.4/1.8* double mutant plants (**Figure 6**). When wild-type pistils were pollinated with *ltp1.4/1.8* mutant pollen, *in vivo* pollen tube growth was comparable to those of the controls (**Figure 6A** and **6B)**. Conversely, when pistils of the *ltp1.4/1.8* double mutant plants were pollinated with wild-type pollen, defective pollen tube targeting was observed (**Figure 6C**), indicating that the apparent defect in either pollen tube emergence or funicular and micropylar pollen tube guidance seems associated with the female floral organ functionality.

**Figure 6.**
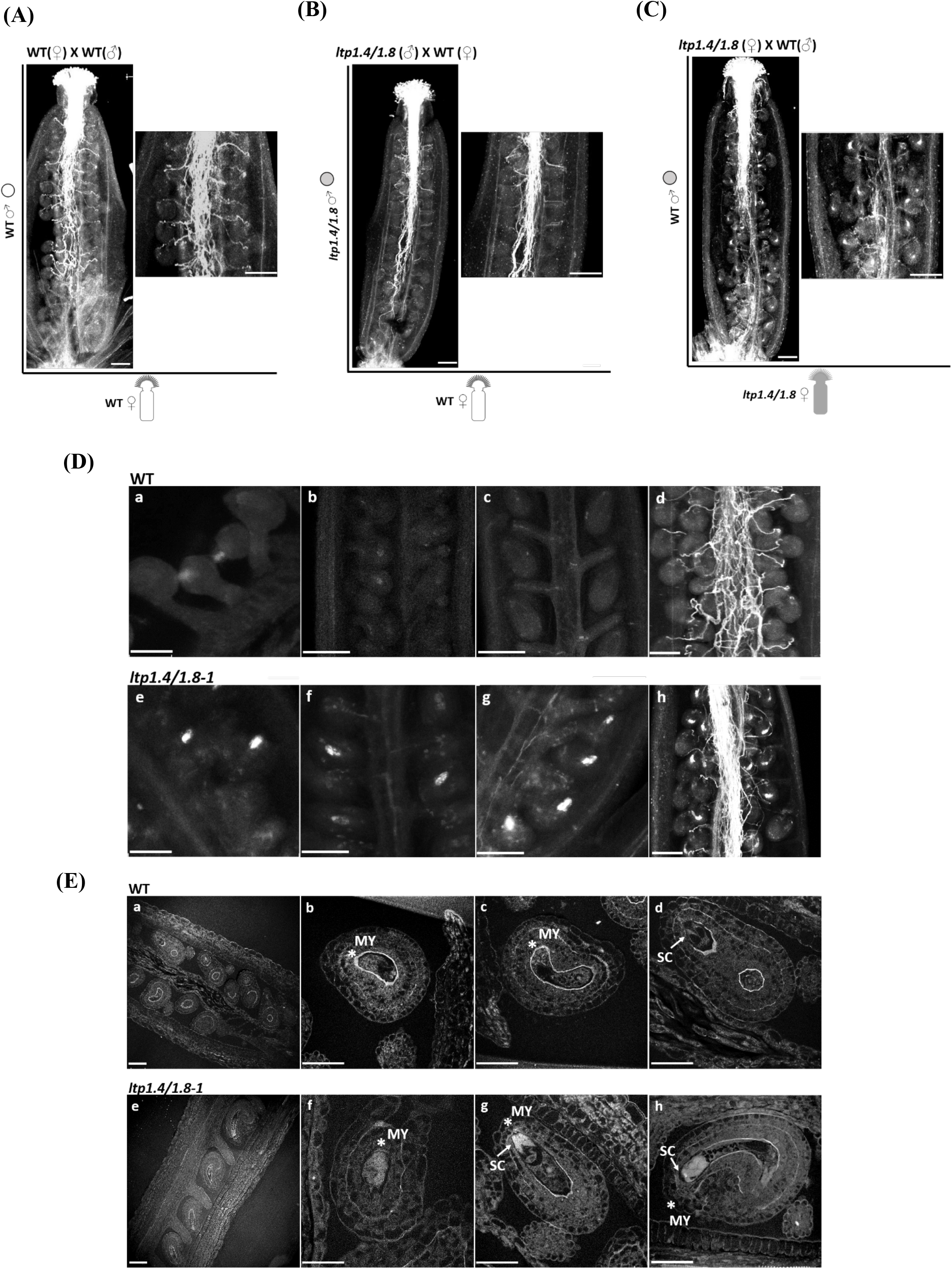
Loss of LTP1.4 LTP1.8 causes a maternal effect seed abortion phenotype. **(A-C)** *In vivo* reciprocal cross-pollination of *ltp1.4/1.8* to the wild-type plants. **(A)** Wild-type pollen as donor and wild-type pistils as acceptor, **(B)** *ltp1.4/1.8* pollen as donor and wild-type pistils as acceptor, **(C)** Wild-type pollen as donor and *ltp1.4/1.8* pistils as acceptor. No callose deposition was observed in the wild-type ovules when pollinated with wild-type **(A)** and mutant pollen **(B)**, but callosized ovules were detected when *ltp1.4/1.8-1* pistils were pollinated with wild-type pollen **(C)**. Aniline-blue staining after 24hrs after pollination (HAP) shows *in vivo* pollen tube growth (n=10); Scale: 100μm. **(D)** Analysis of callose deposition pattern in wildtype and *ltp1.4/1.8-1* mutant ovules in course of development. **(a, b, c, d)** Wild-type ovules during megasporogenesis and megagametogenesis. (a) Visible callose staining around the megaspores. (b) Stage 3-II (two-nuclear embryo sac), faint callose is visible in the young developing ovules. (c) No visible callose deposition at stage 3-V (eight-nuclei embryo sac). (d) No visible callose at postfertilization stage (4-VI), mature fertilized flower. **(e, f, g, h)** Ovule development of *ltp1.4/1.8-1* mutant. (e) Visible callose deposition in the megaspores. (f) Ovule showed fluorescent callose at stage 3-II (two-nuclear embryo sac). (g) Stage 3-V (eight-nuclei embryo sac), increased amount of callose deposition in the ovules. (h) After fertilization, callose deposit still persist in the ovules; Scale: 100μm. **(E)** Overview of callose deposition in transverse sections of Arabidopsis ovules. **(a-d)** In wild-type ovules callose fluorescence is visible only around the nucellus periphery. **(e-f)** In *ltp1.4/1.8-1* ovules, the callose deposition was found to be concentrated around the synergid cells at the micropylar end. Aniline blue staining was performed on the 2μm semi-thin sections of ovules from matured flowers (stage-14) embedded in Technovit-7100 resin; Scale: 100μm. MY: micropylar end, SC: synergid cells.

### *Ltp1.4/1.8* double mutants exhibit aberrant callose deposition at the micropylar region

Closer inspection of the aniline-blue staining revealed aberrant callose depositions in ovules of the double mutant lines and further suggested that the *ltp1.4/1.8* female gametophyte (embryo sac) might be responsible for the observed fertilization phenotype. In order to determine the dynamics of callose depositions more precisely, ovules were dissected at different developmental stages and stained with aniline blue to visualize callose in the developing ovules. In wild-type as well as *ltp1.4/1.8* mutant ovules, callose depositions were visible around developing megaspores already during early stages of embryo sac development (**Figure 6D, a**). At later stages of megagametogenesis (stages 3-V, eight-nuclei embryo sac and 3-VI, mature embryo sac) (Schneitz et al., 1995), callose progressively became absent from wild-type ovules (**Figure 6D, b-c**). In contrast, ovules at stages 3-V and 3-VI of the *ltp1.4/1.8* mutants exhibited sustained, strong fluorescence (**Figure 6D, e-g**). This suggests that in wild-type, following establishment of the functional megaspore, initial callose depositions are gradually degraded during the process of embryo sac development to entirely disappear upon embryo-sac maturation. Likewise, callose was not observed in the fertilized ovules of wild-type plants (**Figure 6D, d**). In the *ltp1.4/1.8* self-fertilized plants the presence of callose was still visible after pollination (**Figure 6D, g-h**) and high levels of callose remained in the central region of the embryo sac during the late development of female-sterile *ltp1.4/1.8* ovules in comparison to wild-type.

To localize callose deposition in ovules more precisely, we performed callose staining on 2 μm semi-thin sections of ovules from Technovit embedded mature flowers (stage-12). Fluorescence microscopy revealed a distinct layer of callose deposition surrounding the embryo sac of wild-type mature ovules, that appeared open towards the micropylar end (**Figure 6E, a-d**). On the contrary, in addition to a thin callose layer surrounding the embryo sac, ovules of the *ltp1.4/1.8* double mutant exhibited prominent callose depositions inside the embryo sac, concentrating around the synergids, primarily at the micropylar end (**Figure 6E, e-h**). Taken together, these results implicate, that AtLTP1.4 together with AtLTP1.8 possess an important role in pollen tube attraction via a yet unknown molecular mechanism regulating callose homeostasis in the developing ovule.

## DISCUSSION

### The diverse expression pattern of Arabidopsis LTPs in the floral tissue suggest a distinct role for certain LTPs in the reproduction process

Plant non-specific lipid transfer proteins, represent a ubiquitous prolamin family of proteins found in all land plants, but absent in charophyte green algae or any other organism (Edstam et al., 2011; Salminen et al., 2016). Despite their suggested role for lipid binding and transport (Zhang et al., 2010; Edqvist et al., 2018) (**Supplemental Figure 4**), *in vivo* functions of LTPs remain rather enigmatic. However, increasing evidence implies a role for certain LTPs in the transfer and deposition of lipid monomers required for assembly of cutin (Salminen et al., 2018) and cuticular wax (Cameron et al., 2006; Kim et al., 2012; Lee and Suh, 2013), suberin (Deeken et al., 2016), and sporopollenin (Zhang et al., 2010) present on many plant surfaces. Thus, LTPs are considered key proteins for colonization of land plants. More recently, it was suggested that some LTPs may be involved in other processes, including plant development or signaling during pathogen attack (Kim et al., 2008; Jülke and Ludwig-Müller, 2016).

Of the 756 CRP-encoding genes present in the Arabidopsis genome, 205 and 139 CRP genes were reported as specifically expressed in the female and male gametophyte, respectively (Huang et al., 2015). Thus, about half of all Arabidopsis CRP genes are involved in gametophyte development and/or reproduction, including LTPs. The latter hypothesis is corroborated by our finding that certain LTPs (such as (*AtLTP1.5, AtLTP1.4, AtLTP1.12, AtLTP1.11, AtLTP1.8, AtLTP1.6* and *AtLTP1.7*) are abundantly expressed in floral tissues.

GUS-reporter-mediated localization and expression analysis of these six members of the *type-I nsLTP* genes revealed distinct but partially overlapping gene expression patterns during seedling growth (see **Supplemental Figure 1**) and in particular during flower development (**Figure 1**). In the Arabidopsis flower, only *AtLTP1.4* and *AtLTP1.8* were strongly expressed in stigma papillar cells during the young bud stage of the mature flowers, suggestive for their plausible role in the synthesis and secretion of the stigmatic exudate to facilitate pollen adhesion and/or hydration. Moreover, *AtLTP1.4* and *AtLTP1.8*, *AtLTP1.12*, *AtLTP1.11*, *AtLTP1.6* and *AtLTP1.7* were mutually expressed in the stylar region of the pistil (Figure 2 and Figure 4) where the pollen tube grows in to reach the ovules. Furthermore, we found *AtLTP1.4, AtLTP1.12* and *AtLTP1.8* expressed in primary female reproductive tissues such as the funiculus or ovules (**Figure 2**). In agreement with previous studies (Thoma et al., 1993; Arondel et al., 2000; Tung et al., 2005), our findings let us conclude that depending on their individual expression profiles – these LTPs may exert their function in providing competency during the pollen tube growth and guidance along the transmitting tract (TT) and possibly in ovular guidance. This hypothesis is further supported by our finding that with the exception of AtLTP1.7 – pistil expressed LTPs were transcriptionally induced upon pollination (**Figure 4**). To date, however, except for a *AtLTP1.8* gain-of function mutant (Chae et al., 2009), no-other LTP has been functionally characterized with respect to its function in pollen tube guidance and fertilization success.

Notably, the majority of peptides reported to be involved in the communication and signaling process during fertilization and seed development are secreted as extracellular CRPs. A molecular survey of several reproductive tissues (like pollen and ovary) in *Zea mays* revealed a prominent expression of genes coding for different CRP groups like defensin-like proteins (DEFLs) possibly contributing to the reproduction process (Li et al., 2014). Among them, the egg cell expressed *Zea mays* EGG APPARATUS 1 (ZmEA1) (Dresselhaus et al., 1994) was identified as an attractant peptide for micropylar guidance (Márton et al., 2012) while *Zea mays* EMBRYO SAC 4 (ZmES4) was shown to trigger pollen tube burst and sperm cell release (Amien et al., 2010). In Arabidopsis and Torenia defensin-like CRPs of the LURE family have been shown to represent diffusible signals that mediate micropylar guidance upon binding to their cognate leucine-rich repeats (LRR) receptor-like kinases (Qu et al., 2015; Takeuchi and Higashiyama, 2016). Despite their distinct molecular modes of action these CRPs have in common to be actively secreted into the extracellular spaces outside the plasma membrane – the extracellular matrix (ECM) – to fulfill their signaling role.

### Diverse subcellular localization patterns of Arabidopsis LTPs

The possible signaling function of LTPs might correlate with the existence of a common N-terminal signal peptide. It suggests that LTPs, alike many other CRPs, usually localize to the ECM to exert their possible function in pollen adhesion, directional growth and guidance of pollen tube and seed development. Our studies on the sub-cellular localization of gametophyte expressed LTPs revealed that AtLTP1.4, 1.8, and 1.6 indeed localize to the apoplast (**Figure 2** and **Supplemental Figure 2A, 2E**). Despite the existence of a predicted N-terminal signal peptide however, AtLTP1.7, 1.11, and 1.12 were associated with the plasma membrane (**Supplemental Figure 2B, 2D, 2E**). In contrast to G type LTPs (Lee et al., 2009), *AtLTP1.7*, *AtLTP1.11* and *At*LTP1.*12* proteins neither harbor a glycosylphosphatidyl anchor (GPI anchor) nor a transmembrane domain. Therefore, the association of these LTPs, as a peripheral protein, with the plasma membrane would probably rely on ionic-protein-lipid interaction and hydrophobic stability. Our finding, that certain LTPs are secreted to the ECM, while others are associated with the plasma membrane, further points to the possibility that some LTPs are secreted via a ‘constitutive’ pathway, while other LTPs take the route of a so far noncharacterized ‘triggered’ secretory pathway.

### LTP1.4/1.8 seem to act in a complementary manner to exert their function in fertilization

Among the floral expressed LTPs, only AtLTP1.8 has been assigned a possible role in fertilization. An *AtLTP1.8* gain-of-function mutant displays abnormal pollen tube guidance and reduced fertilization success, possibly through interaction of AtLTP1.8 with pectin in the pollen tube (Chae et al., 2009). None of the investigated single knock-out mutants for LTPs in this study showed a phenotype related to gamete development or fertilization, which might be explained by existent functional redundancy.

On the other hand, and as reported for CRPs, including LURE and RALF peptides together with their corresponding RLKs receptors (Zhang et al., 2017), LTP action might rely on their functional interaction complexes. Indeed, when analyzing CRISPR/Cas9-generated double mutants of the highly homologous AtLTP1.4 and AtLTP1.8, we observed reduced seed set due to the decreased numbers of fertilized eggs (**Figure 5**). While pollen tube germination and growth along the transmitting tract was wild-type-like in the mutant, our results showed that the concomitant loss of both, AtLTP1.4 and AtLTP1.8 gene functions, caused aberrant callose deposition patterns in the *ltp1.4/1.8* mutant ovules. Callose represents a cell wall β-1,3-glucan polymer and alike for other angiosperms, in wild-type Arabidopsis plants plays an essential role during megasporogenesis (Newbigin et al., 2009; Zhou et al., 2016) of female reproductive development. Typically, callose deposition is first observed in the wall of the megasporocyte, and after meiosis in the walls of the megaspore tetrad (Schneitz et al., 1995). It is hypothesized that the general function of callose during the process of megasporogenesis is to act as a temporary isolation barrier, preventing the meiocyte from cohesion and fusion. In addition, callose might suppress the non-functional megaspores by isolation, thereby ensuring that only the functional megaspore participates in megagametogenesis (Webb and Gunning, 1990). Finally, callose is steadily degraded as the functional megaspore undergoes mitosis, and thus, callose is almost absent from the fully developed embryo sac. This observation is well in line with our results obtained with developing wild-type ovules (**Figure 6**). Based on aniline blue staining, the embryo sac interior of wild-type ovules was essentially free of callose. We observed however, that it was engulfed by a sharp callose layer (**Figure 6E**) that interestingly tended to open towards the micropylar end. The position of this callose boundary appears reminiscent of a recently reported cutin-based apoplastic lipid barrier that is deposited by the ovule inner integument during embryo sac development (Coen et al., 2019), but different from the seed coat cuticle, covering the endosperm outer surface (Demonsais et al., 2020). Thin sections revealed that this sharp callose boundary was also present in the *ltp1.4/1.8* double mutant ovules, but in contrast to wild-type, massive callose accumulation was also apparent inside the embryo sac of the *ltp1.4/1.8* mutants. Remarkably, callose deposition was prominent around the synergid cells at the micropylar end. The massive accumulation of callose at the micropylar end of *ltp1.4/1.8* mutants could result in functional megaspore dysplasia, leading to embryo abortion through inefficient metabolite and nutrient transport. Indeed, callose synthesis represents an early indicator of imminent ovule abortion and has recently been shown to be associated with premature termination of plant reproduction during stress (Sun et al., 2004). Data from *ltp1.4/1.8* self-pollination and cross-pollinations between the *ltp1.4/1.8* mutant and wild-type plants though rather suggest a role of AtLTP1.4 and AtLTP1.8 in pollen tube guidance *in vivo*. In this context, callose depositions at the micropylar end could act as a diffusion barrier for synergid derived pollen tube attractants (Pagnussat et al., 2007). In support of this finding, we observed that pollen tubes largely failed to emerge from the transmitting tract in the *ltp1.4/1.8* mutants (Figure 5E middle and right). Since in Arabidopsis, the distance between the micropyle and the transmitting tract is about100 μm, but LURE-dependent guidance has an effective range of only 20 μm, it is probably not the LURE-dependent guidance cue that is compromised in the *ltp1.4/1.8* mutant but rather so far unknown ovule-derived long-range signals (Mizuta and Higashiyama, 2018).

In our study, we observed sustained deposition of callose in the *ltp1.4/1.8* mutant ovules starting from megaspore formation until the embryo sac maturation. Ovule sterility due to callose deposition, impedes fertilization, thereby contributing to abortion and lower seed set. Our observation that both wild-type and mutant pollen failed to fertilize the callosized ovules in *ltp1.4/1.8* pistils, indicates a possible role of these LTPs in orchestrating the perception and final re-orientation of pollen tube and penetration into the micropyle. These results also explain the reduced seed set observed in the *ltp1.4/1.8* mutant and it indicates that the combined mutation affects callose homeostasis in the female gametophyte, probably by disturbing the homeostasis between its synthesis and degradation process. Callose homeostasis is tightly regulated by the antagonistic action of two types of enzymes, i.e. callose synthases (CalSs) and β-1,3-glucanases (BGs), which respectively confer synthesis and degradation of the β-1,3-glucan polymer.

While CalS proteins represent transmembrane proteins, it is already reported that, GPI-anchored callose binding (CBs) proteins and β-1,3-glucanases (BGs) are transported to the plasma membrane as their target location. In addition, membrane composition appears critical for proper translocation of CBs and BGs and is mediated through a lipid raft vesicle-mediated exocytosis process which regulates callose homeostasis during plasmodesmata symplasmic movement (Amsbury et al., 2017; Liu et al., 2019). It is thus tempting to speculate, that the non-vesicular transport of sterols or complex sphingolipids is also mediated by defined LTPs providing proper targeting of CBs and BGs to the plasma membrane (PM) micro domains i.e., lipid rafts. AtLTP1.4/1.8, present in the apoplast, could act as sterol/sphingolipid carriers modulating PM micro domains at the extracellular leaflet of the PM and their lack could affect PM micro domain lipid composition and thus result in the improper targeting or assembly of signalosomes involved in the regulation of callose homeostasis.

Our findings open doors for the study of specific LTPs in the context of fertilization. Considering the role of LTPs, either as signaling, scaffolding or transport proteins, together with their broad differential expression along male and female reproductive tissues, it is unquestionable that especially the AtLTP1.4/1.8 pair appears as a multifunctional protein moiety. Future work is now required to elucidate the detailed molecular interaction between LTPs and their potential partners to reveal their enigmatic role in callose homeostasis and the pollen tube guidance mechanism.

## MATERIALS AND METHODS

### Plant Materials and Growth Conditions

Wild type seeds (Col-0) and the *ltp* knock-out lines (**Supplemental Table 2**) were obtained from the Arabidopsis stock center (NASC, Loughborough, Leicestershire, UK). All plants were grown in an environmentally controlled growth chamber (150μE m^−2^s^−1^ illumination; 16hrs light/8hrs dark; 22°C; 60% relative humidity). Individual plants (generation T1) from each T-DNA insertion line were genotyped to select individual segregated wild-type, heterozygous, and homozygous individual mutant lines (**Supplemental Table 3**).

### qRT-PCR analysis

Tissue was harvested from plants and placed immediately into liquid nitrogen. To compare the expression of different LTPs in the pollinated and non-pollinated pistils, closed buds of stage 9 (Smyth et al., 1990) were used. For pollinated samples, the emasculated pistils were pollinated with fresh pollen and harvested after 1 day of pollination, whereas for non-pollinated sample, the naked pistils from closed buds were harvested after 2 days of emasculation. To minimize the biological variation, three replicates were collected for each experiment containing 30-40 pistils per replicate. Total RNAs were extracted using the RNeasy Mini Kit (Qiagen). Samples were treated with DNase I (Thermo Scientific) and 2μg RNA were transcribed into cDNA using Moloney murine leukemia virus reverse transcriptase (Promega). A 20-fold dilution of the cDNA reaction was used for PCR amplification with the ABsolute QPCR SYBR Green Capillary Mix (Thermo Scientific) in a LightCycler (Roche). ACT2/8 (AT3G18780/AT1G49240) were used as internal standards to normalize for variation in the amount of cDNA template. The PCR program consisted of a first step of denaturation and Taq activation (95°C for 5mins) followed by 45 cycles of denaturation (95°C for 15s), annealing (60°C for 15s), and extension (72°C for 10s). To determine the specificity of the PCR, the amplified products were subjected to melt curve analysis using the machine’s standard method. Transcript numbers were calculated according to the protocol of Szyroki et al., 2001. For each gene, four reactions were carried out, including two technical replicates and two biological replicates using the primers listed in (**Supplemental Table 4**).

### *In vivo / In vitro* pollen tube growth assay

All *in vitro* pollen tube experiments were performed as described by Boavida and McCormick, 2007. Flowers from *Arabidopsis thaliana* plants two weeks after bolting were used as source of pollen. Pollen grains were germinated on a solid germination medium 1mM CaCl2•2H_2_O, 0.01% (w/v) H_3_BO_3_, 1mM CaCl_2_, 1mM MgSO_4_•7H_2_O, 18% (w/v) sucrose, low melting agar pH 7.5. The freshly opened *Arabidopsis thaliana* flowers were removed with a pair of tweezers, and the pollen grains were tapped on pollen growth medium and incubated for 6hrs at RT in black box with wet paper used to keep constant saturated humidity for germination. After 6hrs, pollen tube length and morphology were examined using the Keyence microscope. The viability of pollen was calculated based on the germinated and non-germinated pollen.

To view pollen tubes growing inside transmitting tract, emasculated pistils were hand-pollinated with the fresh pollen grains. 12-14hrs later, the pistils were cut and fixed in 10% acetic acid-EtOH (3:1) solution overnight. Tissues were softened in 8M NaOH overnight, washed with 50mM K2HPO4 buffer, pH 7.5, and stained in 0.01% aniline blue. Fluorescent images were observed under confocal laser scanning microscopy (TCS SP5 2; Leica).

To evaluate the fertilization efficiency, mature siliques were measured for their lengths and dissected to identify unfertilized ovules. Siliques from the primary inflorescences of wild type and mutant plants were selected from top to bottom and bleached in 70% ethanol for 48-72hrs. The ethanol-bleached siliques were then imaged under the dissecting microscope to examine the seed set. For *in vivo* reciprocal cross-pollination, 10 floral buds at stage-9 (Smyth et al., 1990) were emasculated per cross a day before hand pollination. Fresh pollen at flower stage 13 were fully applied to the stigma of the emasculated pistil. After 14hrs pollination, the pollinated pistil was fixed, stained with aniline blue, and examined as described above. To visualize the callose deposition in the ovules, mature flowers of stage 13 were used for aniline blue-staining.

### GUS reporter assay

The promoter region (~1 kb) of LTP1.4, LTP1.12, LTP1.11, LTP1.8, LTP1.6, LTP1.7 was amplified from *Arabidopsis thaliana* genomic DNA using the gene-specific primers as shown in (**Supplemental Table 5**). The amplified DNA fragments of the individual LTP promoters were fused with the reporter gene β-Glucuronidase (GUS) and cloned in the pMDC164 binary vector (Curtis and Grossniklaus, 2003) using gateway cloning. The constructed binary vectors were introduced into wild-type *Arabidopsis thaliana* plants by Agrobacterium-mediated transformation using floral dip method (Clough and Bent, 1998). Three independent homozygous transgenic plants were used for the LTP-GUS analysis. Flowers of different developmental stage (closed bud, mature flower, siliques) and 10-days old seedling were incubated in GUS staining solution containing 100mM sodium phosphate (pH 7.0), 10mM Na2EDTA, 1mM XGluc, 0.5mM potassium ferrocyanide, 0.5mM potassium ferricyanide, and 0.1% (v/v) Triton X-100 at 37°C for 2:30hrs. The chlorophyll from stained tissues was removed by incubation in 70% (v/v) ethanol at room temperature overnight. The samples were examined by means of a digital microscope (Keyence, Osaka, Japan).

### *In planta* sub-cellular localization of *At*LTPs by transient transformation of *N. benthamiana*

To generate the *At*LTP-YFP construct, full-length cDNA of AtLTP1.4, AtLTP1.12, AtLTP1.11, AtLTP1.8, AtLTP1.6 and AtLTP1.7 were amplified by PCR using the primers listed in (**Supplemental Table 6**). The *At*LTP cDNA was cloned into binary vector pCAMBIA 2300 (35S:USER:YFP:35STerm) using the advanced uracil excision-based cloning (USER) technique (Jørgensen et al., 2017).

*Agrobacterium tumefaciens* cells of strain GV3101 (Koncz and Schell, 1986) were used for transformation with various constructs and were grown overnight in 5ml Yeast Extract Broth (YEB) liquid medium at 28°C under shaking until OD600 reached 0.8-1. The cells were centrifuged for 5mins at 4000xg and the obtained pellet was re-suspended in agromix (10mM MES-KOH pH 5.6, 0.15mM acetosyringone, 10mM MgCl2) until OD600 reached 1, and incubated for 2hrs at room temperature. *Nicotiana benthamiana* leaves of 3-week-old plants were infiltrated at their abaxial side with the bacterial suspension. In the case of co-infiltration of two bacterial suspensions carrying different constructs, a ratio of 2:1 was used. The DsRED-labelled REMORIN 1.3 (35S:DsRED-REMORIN 1.3) (Demir et al., 2013) and mCherry-labelled LTP1.5 (35SLTP1.5::mCherry) (Deeken et al., 2016) were used as plasma membrane and cell-wall marker respectively. Plasmolysis assay was performed by incubating leaf sections in 0.5M KNO3 for 5-10mins at room temperature. The plants were incubated for 48hrs in a growth chamber before microscopy was performed. An upright confocal laser scanning microscope (TCS SP5 2; Leica) using a 40x apochromatic water immersion lens was used to visualized YFP fluorescence (excitation: 488nm; emission: 514–551nm), RFP fluorescence (excitation: 561nm; emission: 582–622nm), DsRED fluorescence (excitation: 561nm, emission: 560-600nm). Overlay images (mCherry/mVenus) were created using the software LAS AF (Leica Application Suite Advanced Fluorescence 2.4.1; Leica).

### Generation of *ltp1.4ltp1.8* double knock-out Arabidopsis plants using the CRISPR-Cas9 technique

In general, sgRNAs were selected for specificity using CRISPROR (http://crispor.tefor.net/crispor.py), considering the predicted off-target efficiencies. CRISPR/Cas9 constructs were cloned as previously described in Wang et al., 2015. Briefly, for each guide sequence, a set of two complementary oligos designed with Bsa1 cutting sites were annealed and ligated with the *BsaI*-linearized pHEEE401E destination vector through golden gate reaction. The positive transformants were confirmed by colony PCR using U6-26p-FP and U6-29p-RP primers, and later confirmed by sanger sequencing using U6-26p-FP and U6-29p-FP (**Supplementary Table 7**). The final CRISPR/Cas9 binary vector pHEEE401E which includes an egg cell specific promoter (*EC1.2*) was used for the transformation of *Arabidopsis thaliana* Col-0 wildtype plants. Hygromycin resistance plants of the T1 generation were tested for the insertion of the sgRNA-Cas9-T-DNA by PCR with gene specific primers LTP1.4-sgRNA-153 FP/RP and LTP1.8-sgRNA-154 FP/RP, respectively. Targeted gene mutations were detected by sequencing of these PCR products. Plants with an identified LTP1.4 and LTP1.8 double mutation were allowed to self-fertilize and homozygous plants in the T2 generation were detected by sequencing of LTP1.4 and LTP1.8 PCR products.

Based on sanger sequencing, four different double mutant transgenic lines (#P9-P2-P2, #P9-P3-P3, #*ltp1.4ltp1.8-1* and #*ltp1.4ltp1.8-2*) were chosen for constructing next generation sequencing libraries following the manufacture’s protocol (NEB Next Ultra II DNA kit). Sequencing was carried out using 2 × 150 paired-end NextSeq500 (1-2 Mio reads for all samples together) in Core Unit SysMed, IMIB, University of Wuerzburg. Data processing and statistical analysis was performed using CRISPresso computational tool (http://crispresso.rocks/).

### Callose staining of embedded Arabidopsis flowers

Flowers from the wild type and *ltp1.4ltp1.8* were emasculated at stage 12 (Smyth et al., 1990) At 24hrs after emasculation, pistils were collected and fixed using FAA solution (Ethanol 95%, Glacial Acetic Acid 37%, Formaldehyde, H2O). After fixation, the fixed tissue was subsequently dehydrated with different EtOH concentration (50%, 60%, 70%, 85%, 95% 100%, 30mins). The dehydrated tissue was cleared in different concentration of infiltration solution and ethanol solution. The 100% Technovit solution was replaced with infiltration solution (100% Technovit “basis solution + Härter I) and incubated overnight at RT. The infiltrated tissue was transferred to polymerization resin (Infiltration solution + Härter II) and carefully placed inside the gelatin capsule and incubated overnight at RT. After polymerization, the Technovit-embedded flower samples were trimmed using sharp razor blade under dissecting light microscopy. The cross section of 2-4μm was cut with a glass knife using the ultramicrotome Leica EM UC7 (Leica Microsystems). The semi-thin sections were stained for 30mins in 0.1% aniline blue in 0.1M K3PO4 at RT. Slides were overlaid with coverslips and sealed with 1-2 drops of aniline blue. The stained sections were observed within 1hr to detect callose deposition using confocal laser scanning microscopy (TCS SP5 2; Leica).

### Correlative Light and Electron Microscopy

Tobacco plants were processed by following steps to localize the 35SLTP1.8::YFP construct by fluorescence microscopy in its ultra-structural context of electron micrographs. Using a 2mm punch, small discs were punched out of leaves of transfected tobacco and transferred into 200μm deep and 3mm wide high-pressure freezing planchets which were filled with hexadecene. Samples were high-pressure frozen with a Leica EM HPM100 machine at >2100bar pressure and a freezing rate of >20,000K/s and transferred in liquid nitrogen onto frozen solution of 0.1% KMnO4 in acetone. The container was put into a Leica EM AFS2 machine keeping a constant temperature of −90°C and then a freeze substitution process was accomplished according to a previously published CLEM protocol with slight modifications (Markert et al., 2017). In short, a two-component LR-White Medium Grade Acrylic Resin (London Resin Company Ltd.) was used. Samples were polymerized at 40°C for 48hrs. 100nm ultra-thin sections were prepared and transferred onto poly-L-lysine coated glass slides (Thermo Fisher Scientific) with a Leica EM UC7 / FC7 ultracut equipped with a Histo Jumbo Diamond Knife (Diatome). Sections were treated with primary antibody anti-GFP (polyclonal chicken, abcam; product number: ab13970; dilution 1:500 in blocking solution) for 1hr and secondary antibody anti-chicken (goat IgG (H+L) Alexa Fluor 488 conjugate, Thermo Fisher Scientific; 1:500 in blocking solution) for 30mins. Sections were treated with calcofluor-white (0.01% solution in water and mixed with 10% KOH, SIGMA) for 10mins. Sections were imaged by z-stack acquisition with a SIM Zeiss ELYRA S.1. Sections were contrasted with 2.5% uranyl acetate in ethanol for 15mins and with Reynolds’ lead citrate (REYNOLDS, 1963) in H2O (1:1) for 10mins. Sections were mounted on SEM holder and coated with a 2.5nm carbon layer with a CCU-010 Compact Coating Unit machine (safematic). Sections were imaged with a scanning electron microscope JEOL JSM-7500F at 5kV using a *low-angle-backscatter* (LABE) detector. Each selected region of interest (ROIs) were imaged as tile-series manually with a working distance of 5.9-6.0mm and probe current of 0.3nA.

To start the image processing, SIM image stacks were calculated to super-resolution in automatic mode with ZEN black edition software (Zeiss). The image in the focus plane of each SIM image z-stack was extracted. SEM images tiles were stitched with TrakEM2 (Cardona et al., 2012) applying the montage mode *least squares* (*linear feature correspondences*) with a rigid transformation. Image contrast was adjusted with TrakEM2 image filters (1. *Default Min And Max*; 2. *Normalize Local Contrast* (brx/y with 400); 3. *Enhance Contrast*; 4. *Invert*;) and then the montage was repeated. The *blending* function was applied to the images. The SIM image was correlated to the respective SEM image regarding the calcofluor-white signal and cell-wall structure as landmark. Correlation was performed with Inkscape (version 0.91; http://www.inkscape.org) following in principle the previously published CLEM protocol (Markert et al., 2017) and images were finally composed with image editing software GIMP (http://www.gimp.org). For image composing, black pixels of both fluorescence images were converted into transparent and then their layer modus changed to *addition*.

## ACCESSION NUMBERS

The gene sequences mentioned in this study can be found in the GenBank/EMBL data libraries under the following accession numbers: ACTIN2/8 (AT3G18780/AT1G49240), AtLtp1.5 (AT2G38540), AtLtp1.4 (AT2G38530), AtLtp1.6 (AT3G08770), AtLtp1.7 (AT3G51590), AtLtp1.8 (AT3G51600), AtLtp1.11 (AT5G59310) and AtLtp1.12 (AT5G59320).

## AUTHOR CONTRIBUTIONS

D.B., R.D., C.W., C.S., W.D.L., and K.K. conceived and designed the experiments. K.K., M.Z., and S.B. performed the experiments. K.K. and D.B. analyzed the data. D.B., R.D., and K.K. wrote the paper. The authors declare no conflicts of interest that might be perceived as affecting the objectivity of this work.

## Supplemental Information

### Supplemental Data

Supplemental Figure 1. GUS Expression of *AtLTP1.4, AtLTP1.12, AtLTP1.11, AtLTP1.8, AtLTP1.6, AtLTP1.7* in 10 days old Arabidopsis seedling.

Supplemental Figure 2. Sub-cellular localization of AtLTP1.12, AtLTP1.11, AtLTP1.6 and AtLTP1.7.

Supplemental Figure 3. Gene expression levels in T-DNA insertion lines of Arabidopsis nsLTPs and seed set analysis of *ltp* mutant plants.

Supplemental Figure 4. Characterization of Arabidopsis nsLTPs.

Supplemental Figure 5. Mutation analysis of *ltp1.4/1.8-1* and *ltp1.4/1.8-2* plants.

Supplemental Table 1. Comparative analysis of tissue-specific expression of Type I and Type III nsLTPs by using different gene expression databases.

Supplemental Table 2. List of T-DNA insertion lines of Arabidopsis nsLTPs.

Supplemental Table 3. List of gene-specific primer used for genotyping T-DNA insertion mutants.

Supplemental Table 4. List of qPCR primers.

Supplemental Table 5. List of primers used for amplifying LTP promoter.

Supplemental Table 6. List of primers used for amplifying LTP cDNA.

Supplemental Table 7. List of primers used for generating LTP1.4sgRNA and LTP1.8sgRNA.

**Supplemental Figure 1.**
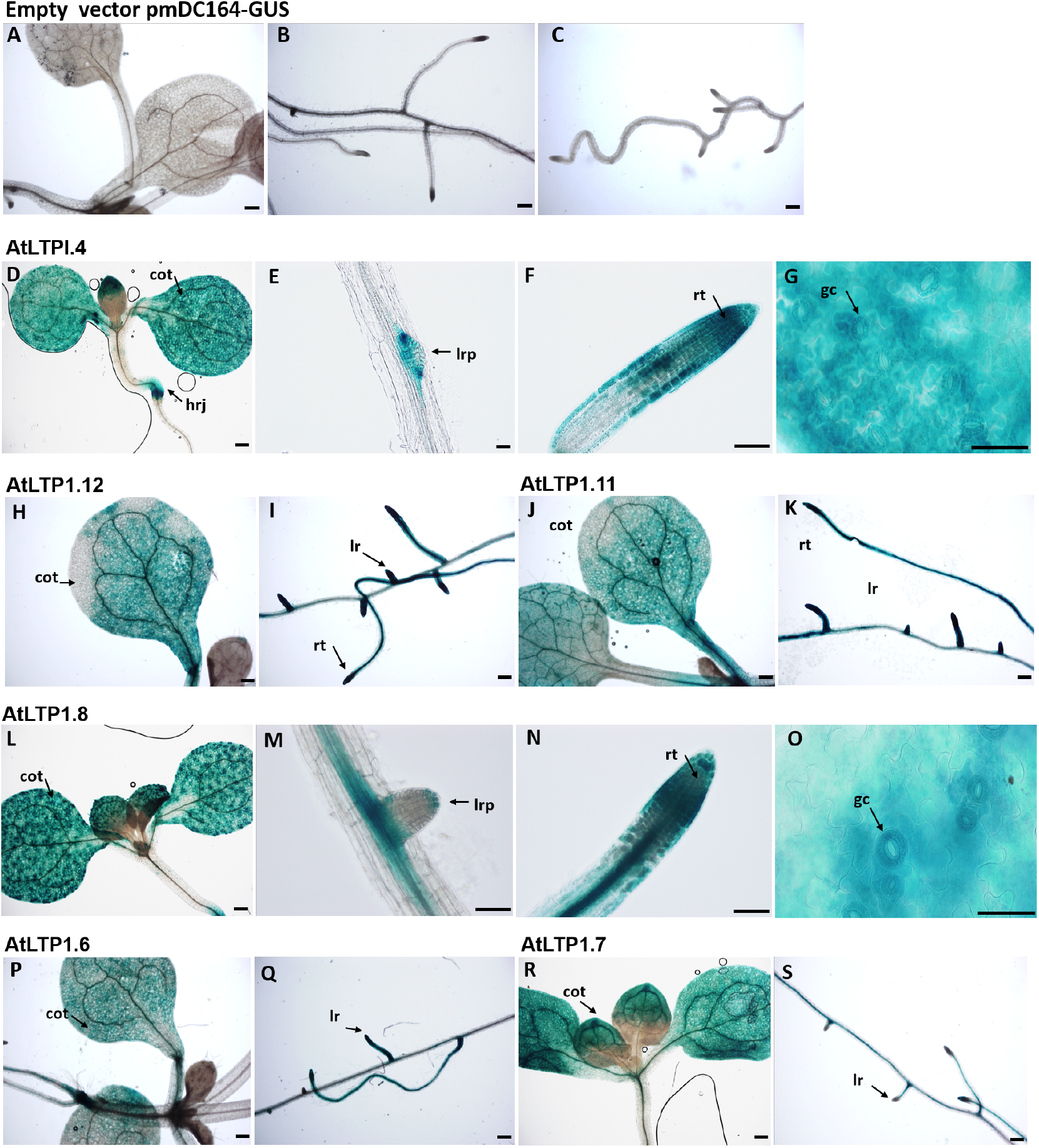
GUS Expression of *AtLTP1.4, AtLTP1.12, AtLTP1.11, AtLTP1.8, AtLTP1.6, AtLTP1.7* in 10 days old Arabidopsis seedling. Histochemical localization of GUS activity in *PromLTP::GUS* plants. (A-C) control, (D-G) *PromLTP1.4::GUS* was observed in the cotyledon, emerging lateral root primordia, primary root tip, vascular bundles and in the guard cells. (H-I) *PromLTP1.12::GUS* and (J-K) *PromLTP1.11::GUS* expression was observed in the cotyledon, root tip emerging and mature lateral root. (L-O) *PromLTP1.8::GUS* expression was localized in the cotyledon, emerging lateral root primordia, primary root tip and in the guard cells. (P-Q) *PromLTP1.6::GUS* and (R-S) *PromLTP1.7::GUS* expression was also detected in the cotyledon and lateral root. Plants were grown under long-day condition (16hrs light/ 8hrs dark). GUS signals were developed for 2:30hrs. cot: cotyledons, hrj: hypocotyl-root junction, lrp: lateral root primordia, rt: root tip, gc: guard cells, lr: lateral root; Scale: 50μm.

**Supplemental Figure 2.**
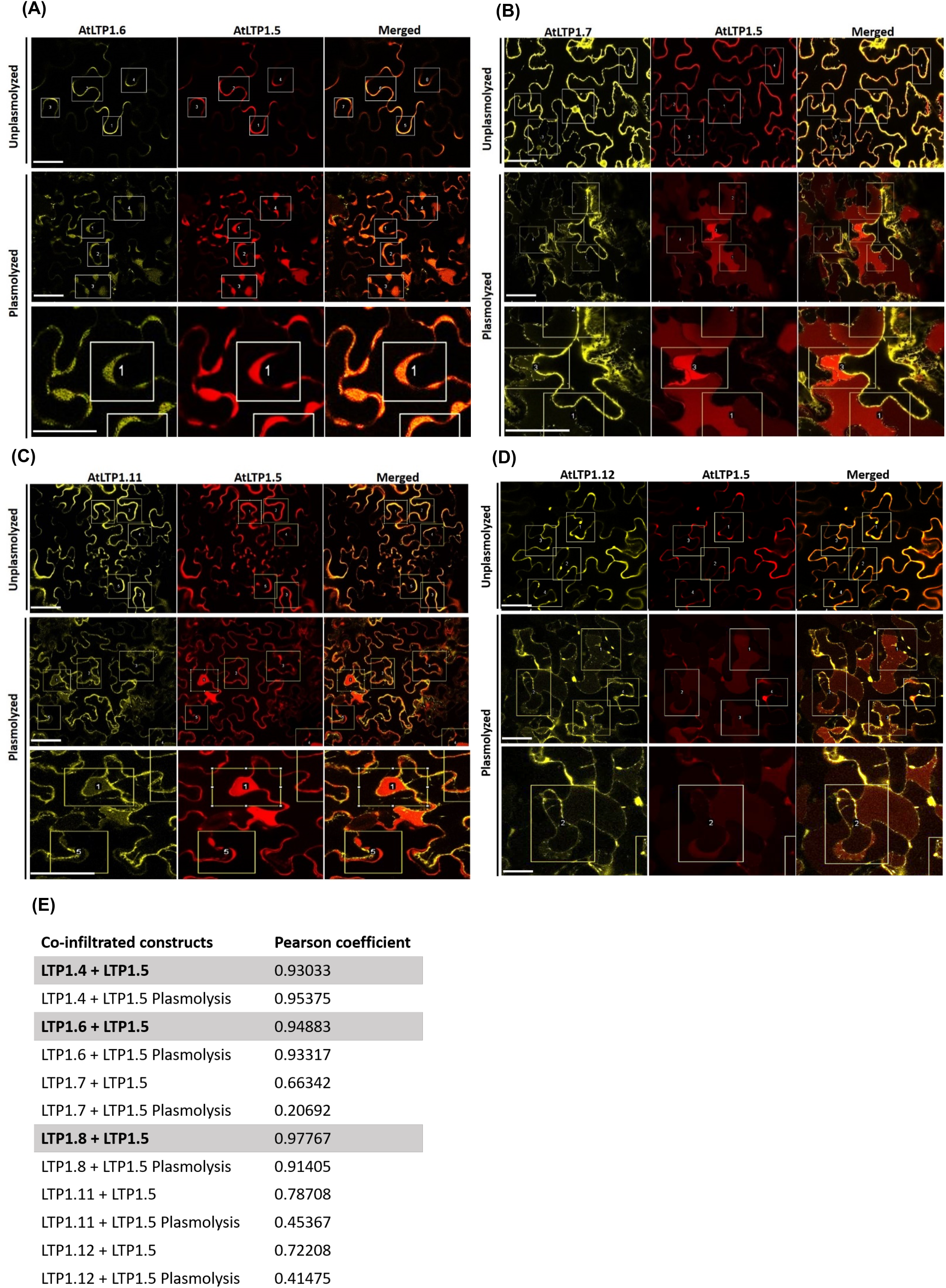
Sub-cellular localization of AtLTP1.12, AtLTP1.11, AtLTP1.6 and AtLTP1.7. **(A)** AtLTP1.6 **(B)** AtLTP1.7 **(C)** AtLTP1.11 **(D)** AtLTP1.12 fused to YFP in *N. benthamiana* epidermal cells imaged 4 days post infiltration. Co-expression of the two-fusion proteins 35SLTP1.6::YFP, 35SLTP1.7::YFP, 35SLTP1.11::YFP, 35SLTP1.12::YFP and cell wall marker 35SLTP1.5::mCherry (upper row). Co-infiltrated leaves were plasmolyzed with 0.5M KNO_3_and documented after 5mins of incubation (middle row). Zoom-in images in the lower row of **(A)** AtLTP1.6 was colocalized with AtLTP1.5 in the apoplastic space (AS) of the plasmolyzed cell as shown in the zoom-in images in the lower row. **(B)** AtLTP1.7, **(C)** AtLTP1.11 and **(D)** AtLTP1.12 shows localization in the retracting plasma membrane (PM) while AtLTP1.5::mCherry is localized in the apoplastic-space (AS) of the plasmolyzed cell. **(E)** The average Pearson’s correlation coefficients ± SEM calculated from 4 randomly selected ROI in each group, n=3. Each selected region of interest (ROI) is marked in yellow box and labelled, respectively; Scale:50 μm. Highlighted LTP1.4, LTP1.8 and LTP1.6 showed higher value of Pearson coefficients for plasmolyzed and non-plasmolyzed cells, suggesting their extracellular localization in the apoplast. The overlap Pearson’s correlation coefficients were calculated using JACoP: Just Another Co-localization Plugin from ImageJ. mCherry (Excitation: 561 nm, Emission: 580– 615 nm; mVenus (Excitation: 514 nm, Emission: 530–555 nm).

**Supplemental Figure 3.**
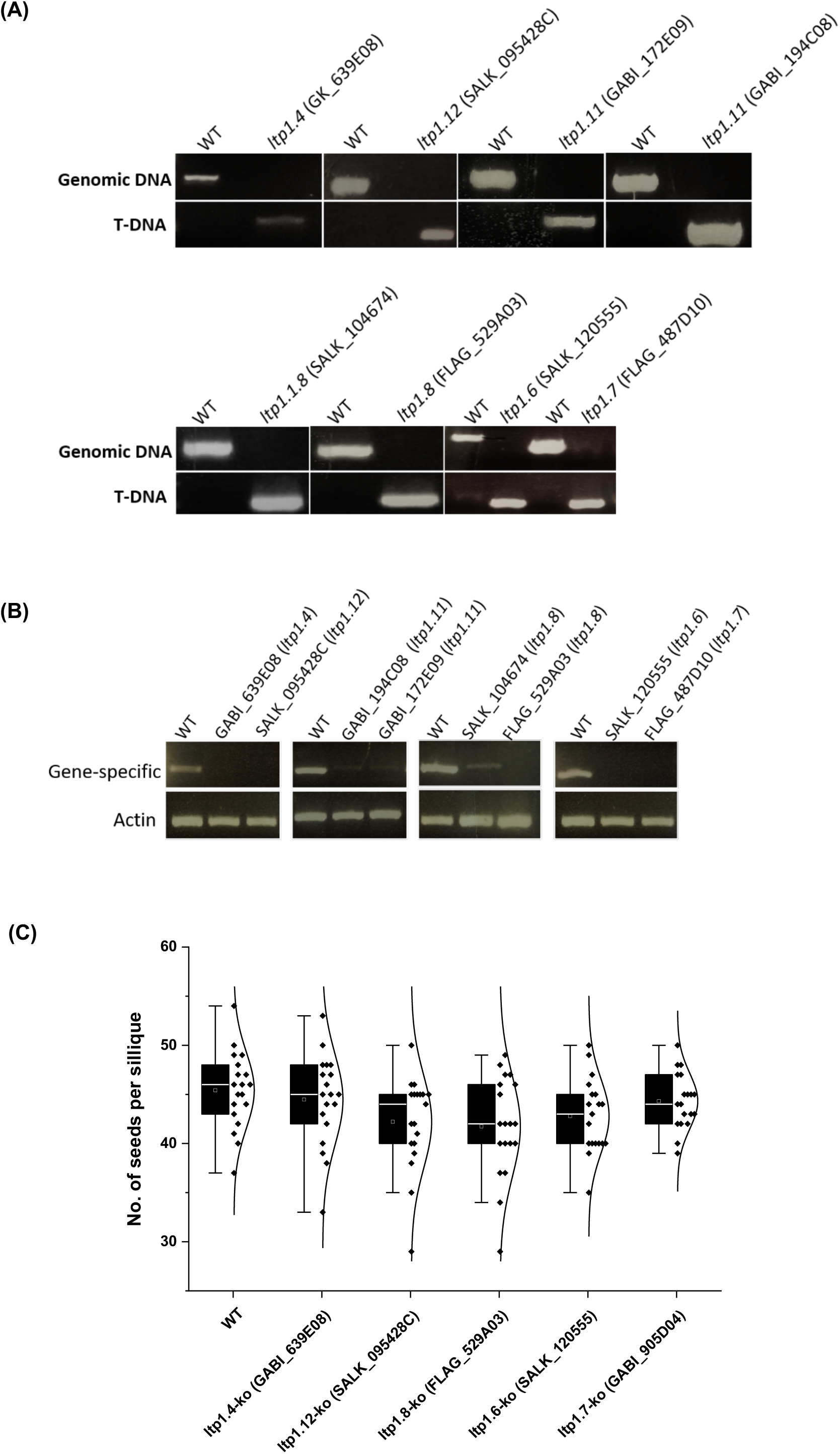
Gene expression levels in T-DNA insertion lines of Arabidopsis nsLTPs and seed set analysis of *ltp* mutant plants. **(A)** Genotyping of LTP T-DNA insertion mutants. PCR based genotyping, amplification of wild-type allele using gene-specific primer (LP, RP) and T-DNA using (LP, T-DNA locus specific primer (for GABI-KAT), LBb1.3 (for SALK), Tag3/Rb4 (for FLAG). **(B)** q-PCR gene expression analysis of LTP T-DNA insertion mutants. Homozygous T-DNA alleles for LTPs, screened by PCR-based genotyping, were evaluated for gene expression by RT-PCR analysis using the gene-specific primer sets and actin (*ACT2/8*) as a control. PCRs amplification were performed in 30 cycles for both LTPs and the actin control. All primers used for PCR-based genotyping and qPCR are listed in Table S2, S3. **(C)** Analysis of seed count revealed no significant differences between WT and *ltp1.4* (GABI_639E08), *ltp1.12* (SALK_095428C), *ltp1.8* (FLAG_529A03), *ltp1.6* (SALK_120555) and *ltp1.7* (GABI_905D04) homozygous knockout lines. Data are shown as mean ± SD and n = 20 siliques from each genotype.

**Supplemental Figure 4.**
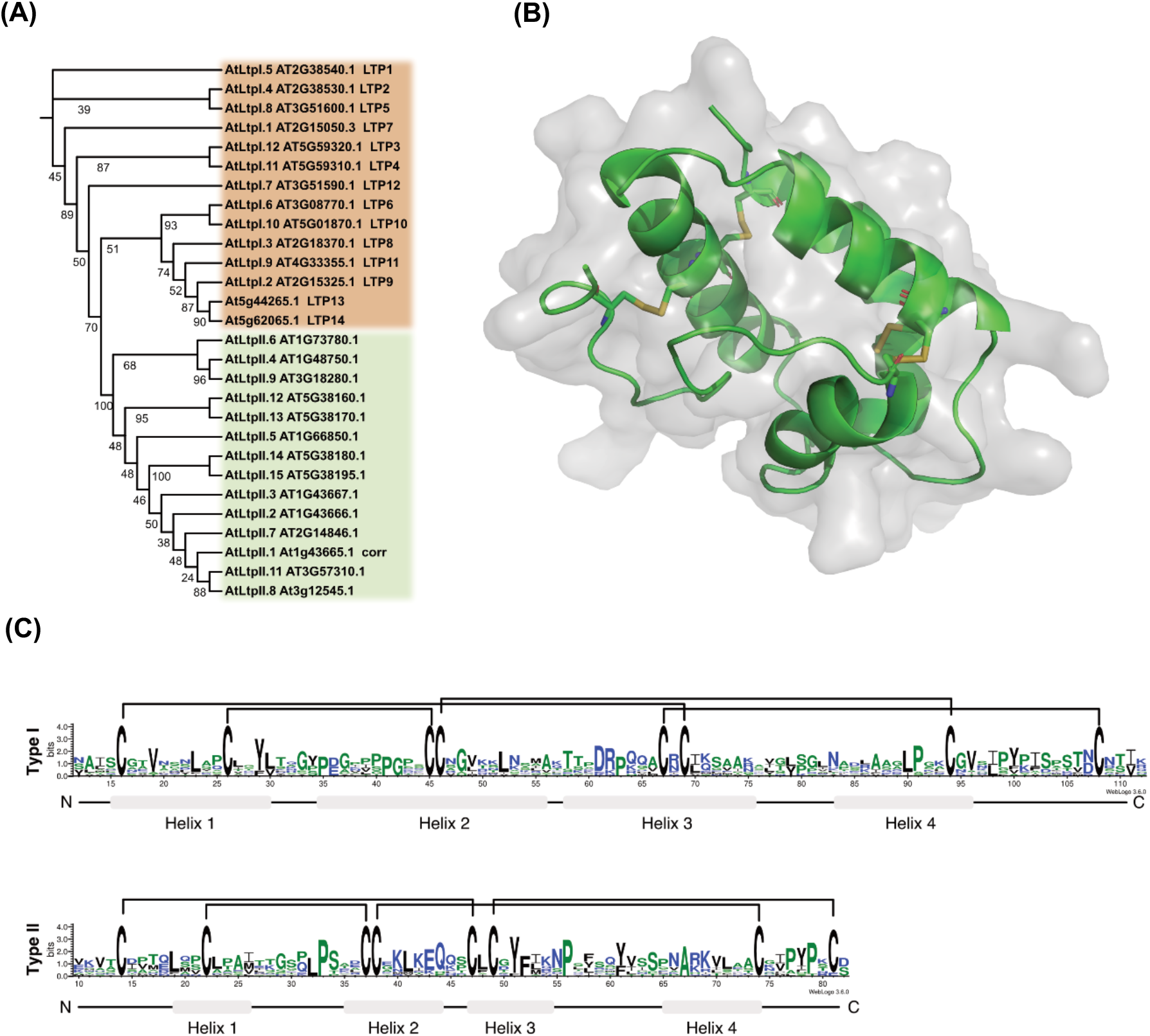
Characterization of Arabidopsis nsLTPs. **(A)** A phylogenetic analysis of Arabidopsis LTPs shows that Type I and Type II LTP proteins separate into distinct clades. Protein alignment was performed on sequences with the signal-peptide removed using MAFFT. Phylogenetic distances were calculated with IQ-Tree using standard parameters with 1000 bootstrap alignments. **(B)** Representation of the 3D structure of Arabidopsis LTP5 by homology modeling with the PyMOL modelling software. **(C)** Amino acid sequence of Type I and Type II nsLTP representing the 8-Cys patterns (primary structure) and four disulfide bridges denoted by the four linkages between cysteine residues. Black linkage depicts the location of four pairs of disulfide bonds indicated by connecting lines in Type I and Type II nsLTPs.

**Supplemental Figure 5.**
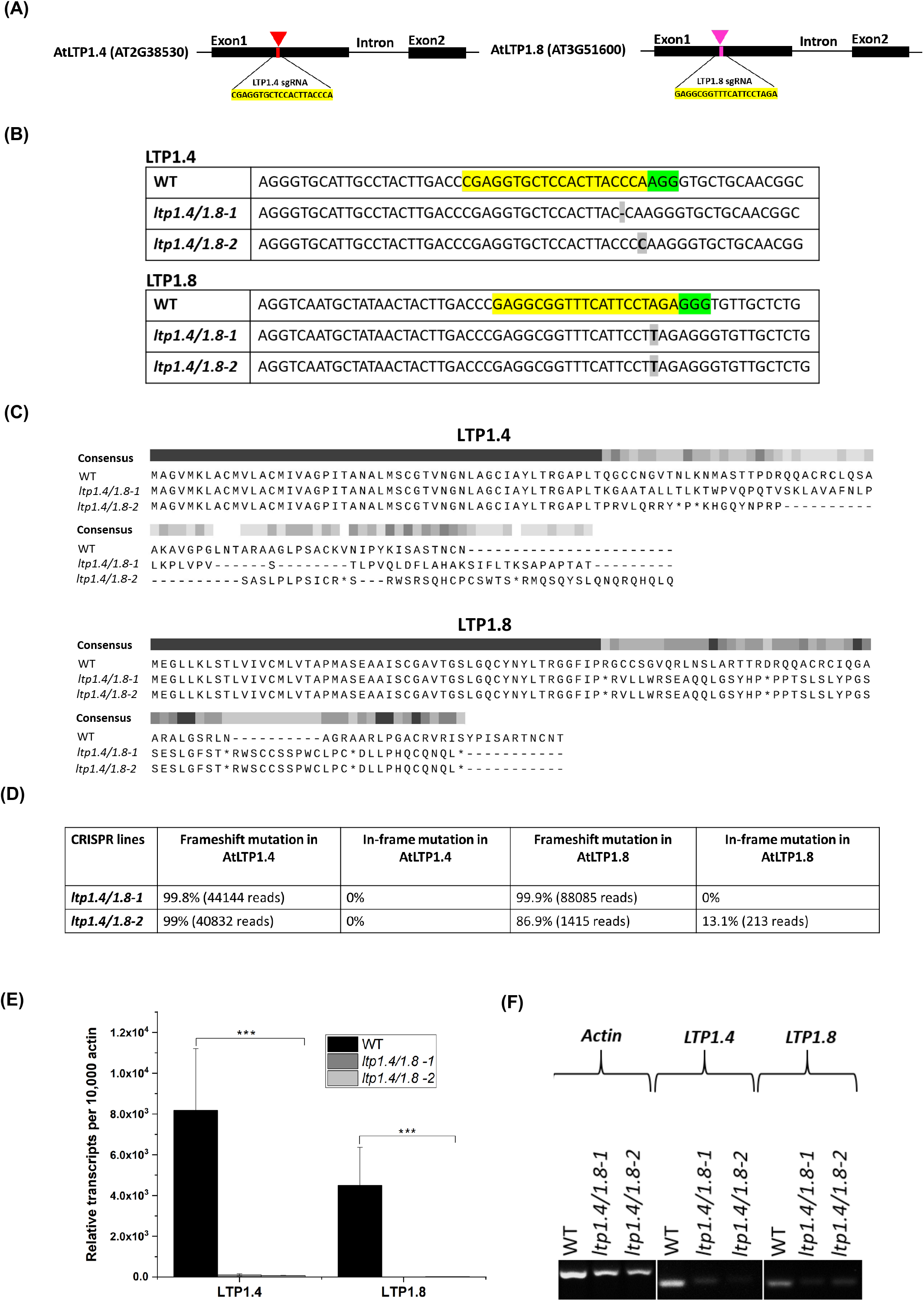
Mutation analysis of ltp1.4/1.8-1 and ltp1.4/1.8-2 plants. **(A)** Scheme showing the targeted site in the AtLTP1.4 and AtLTP1.8 genomic region. The sgRNA^**AtLTP1.4**^ is located in the first exon (132 bp downstream of the start codon), while sgRNA^**AtLTP1.8**^ is located in the first exon (133 bp downstream of the start codon). The sgRNA^**AtLTP1.4**^ and sgRNA^**AtLTP1.8**^ sequence are highlighted in yellow. **(B)** Alignment of T3 homozygous double mutants obtained via EPC CRISPR/cas9 mutagenesis. The targeted region was amplified by PCR and sequenced using standard Sanger sequencing. The sgRNA^AtLTP1.4^ and sgRNA^AtLTP1.8^ sequence are highlighted in yellow and the PAM sequence in green. Deletions and insertions are indicated are highlighted in gr*a*y. **(C)** Comparison of AtLTP1.4 and AtLTP1.8 protein sequences of wild-type and ltp1.4/1.8 mutant lines. Asterisk denotes pre-mature stop codon due to the mutation in the reading frame. **(D)** Analysis of mutagenesis profile from deep sequencing data using CRISPResso2 analysis tool. Quantification of editing frequency as determined by the percentage and number of sequences reads showing frameshift and in-frame mutations. Frameshift and in-frame mutations include any mutations that partially or fully overlap coding sequences, with any non-overlapping mutations classified as noncoding. Unmodified reference reads were excluded from the analysis, and all high dynamic range (HDR) reads were included. **(E-F)** qRT-PCR gene expression analysis of ltp1.4/1.8-1 and ltp1.4/1.8-2 mutants using the gene-specific primer sets (Table S3) and actin (ACT2/8) as a control. PCRs amplification were performed in 35 cycles for both LTPs and the actin control. Data are shown as mean ± SD. The error bars and asterisks indicate values that differ significantly from those of the wild-type plant, n=3; *, *P*<0.05, **, *P*<0.01, ***, *P*<0.001 (*t*-test).

### Supplemental Tables

**Supplemental Table 1.**
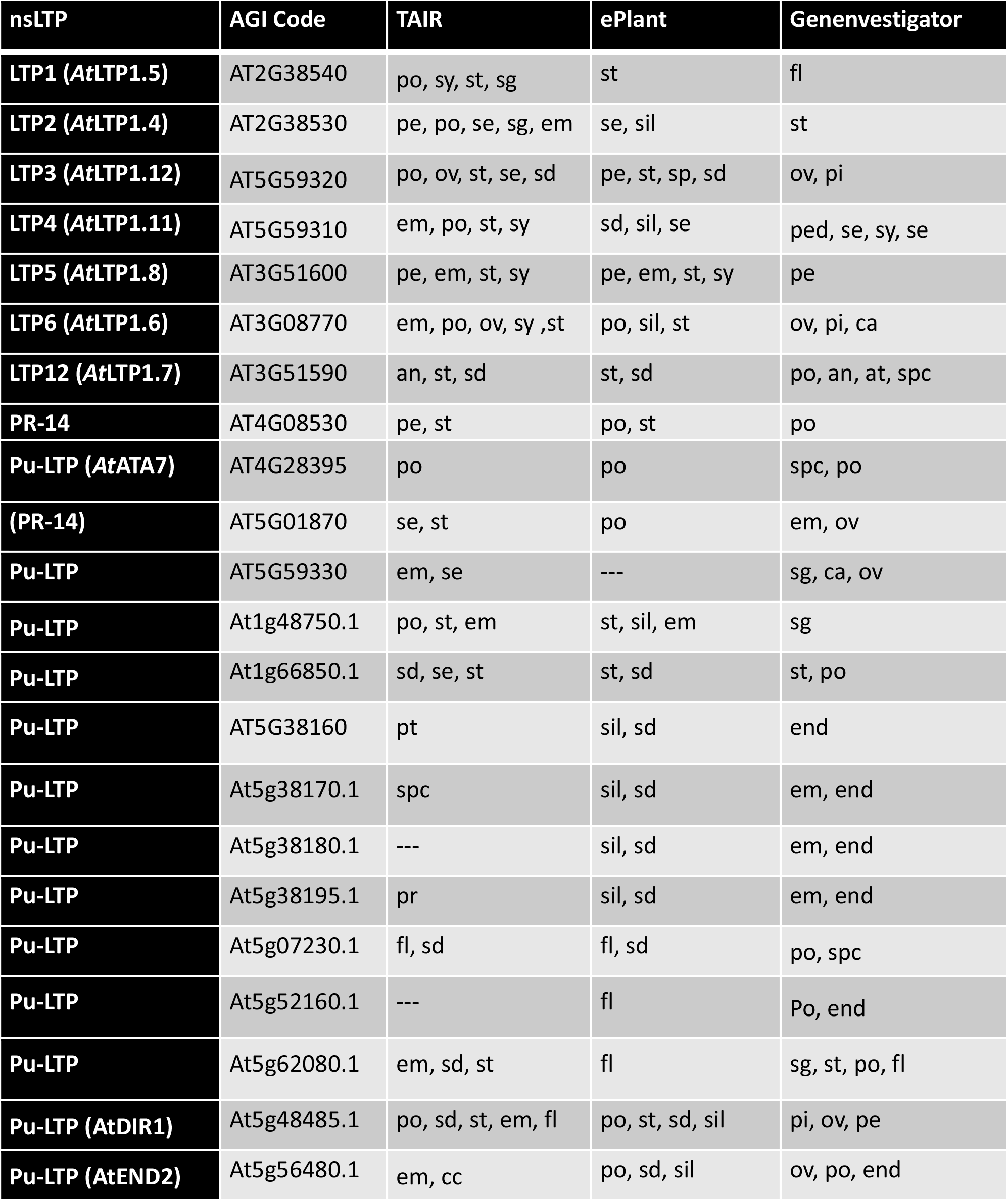
Comparative analysis of tissue-specific expression of Type I and Type III nsLTPs by using different gene expression databases. TAIR (https://www.arabidopsis.org) ePlant (http://bar.utoronto.ca/eplant/), and Genevestigator (https://genevestigator.com) expression database were used for the comparative analysis of tissuespecific expression of individual LTPs in the different part of floral tissue. LTPs lacking information about their expression pattern in the databases were excluded from the analysis. sg-stigma, sy-style, st-stamen, fu-funiculus, ov-ovules, po-pollen, an-anthers, pi-pistil, pt-pollen tube, pe-petals, se-sepals, sd-seeds, em-embryo sac, end-endosperm, fl-different floral developmental stages, spc-sperm cells, sil-siliques, an-anther, pe-petal, pr-primary root, ca-carpel. Pu-LTP: Putative LTP.

**Supplemental Table 2.** List of T-DNA insertion lines of Arabidopsis nsLTPs.

| Gene | AGI code | Accession | T-DNA mutant |
| --- | --- | --- | --- |
| LTP1.4 | AT2G38530 | Col-0 | GABI_639E08-022274 |
| LTP1.12 | AT5G59320 | Col-0 | SALK_095248C |
| LTP1.11 | AT5G59310 | Col-0<br>Col-0 | GABI_194C08<br>GABI_172E09 |
| LTP1.8 | AT3G51600 | WS<br>Col-0 | FLAG_529A03<br>SALK_104674 |
| LTP1.6 | AT3G08770 | Col-0 | SALK_120555 |
| LTP1.7 | AT3G51590 | Col-0<br>WS | GABI_905D04<br>FLAG_487D10 |

**Supplemental Table 3.**
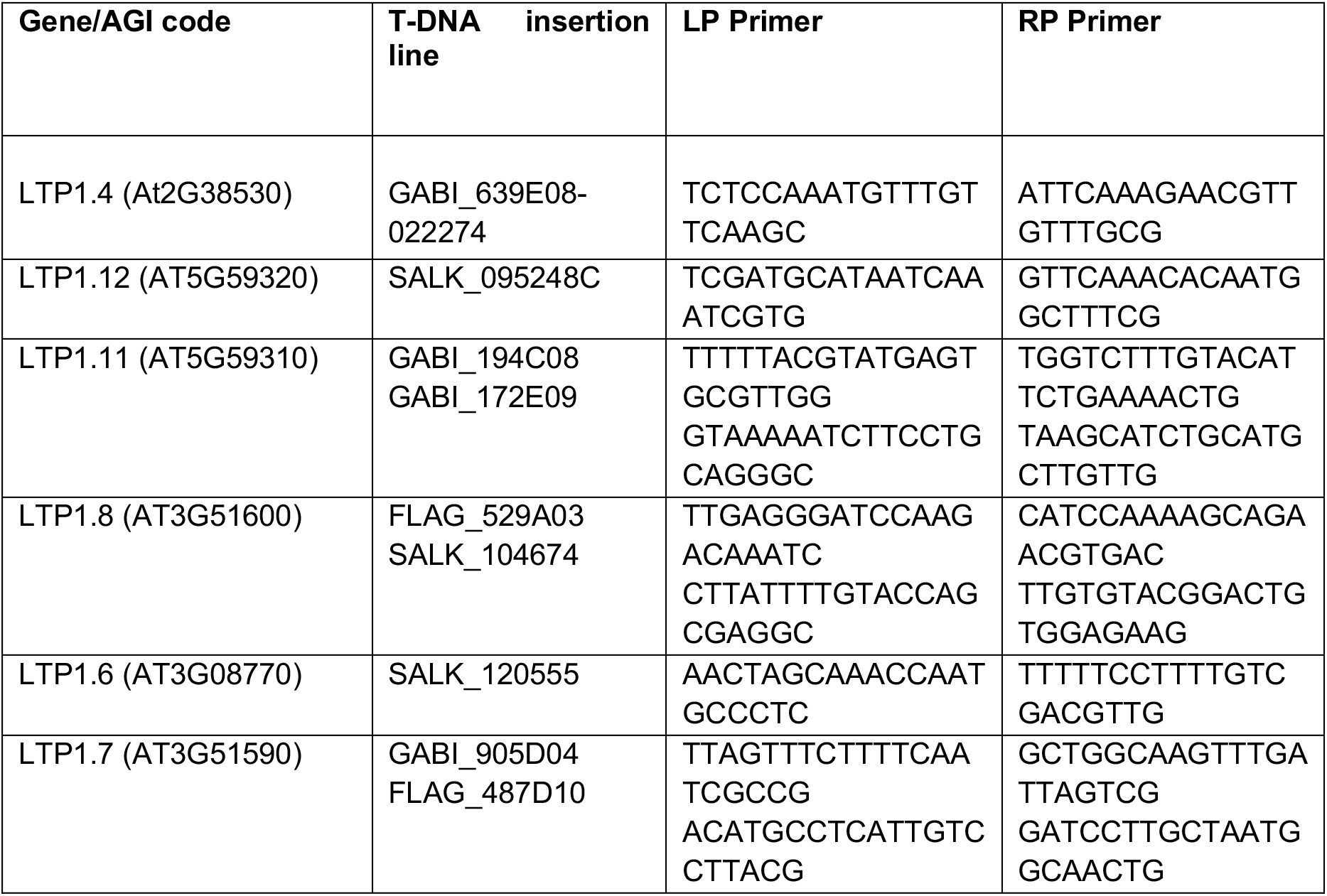
List of gene-specific primer used for genotyping T-DNA insertion mutants.

**Supplemental Table 4.**
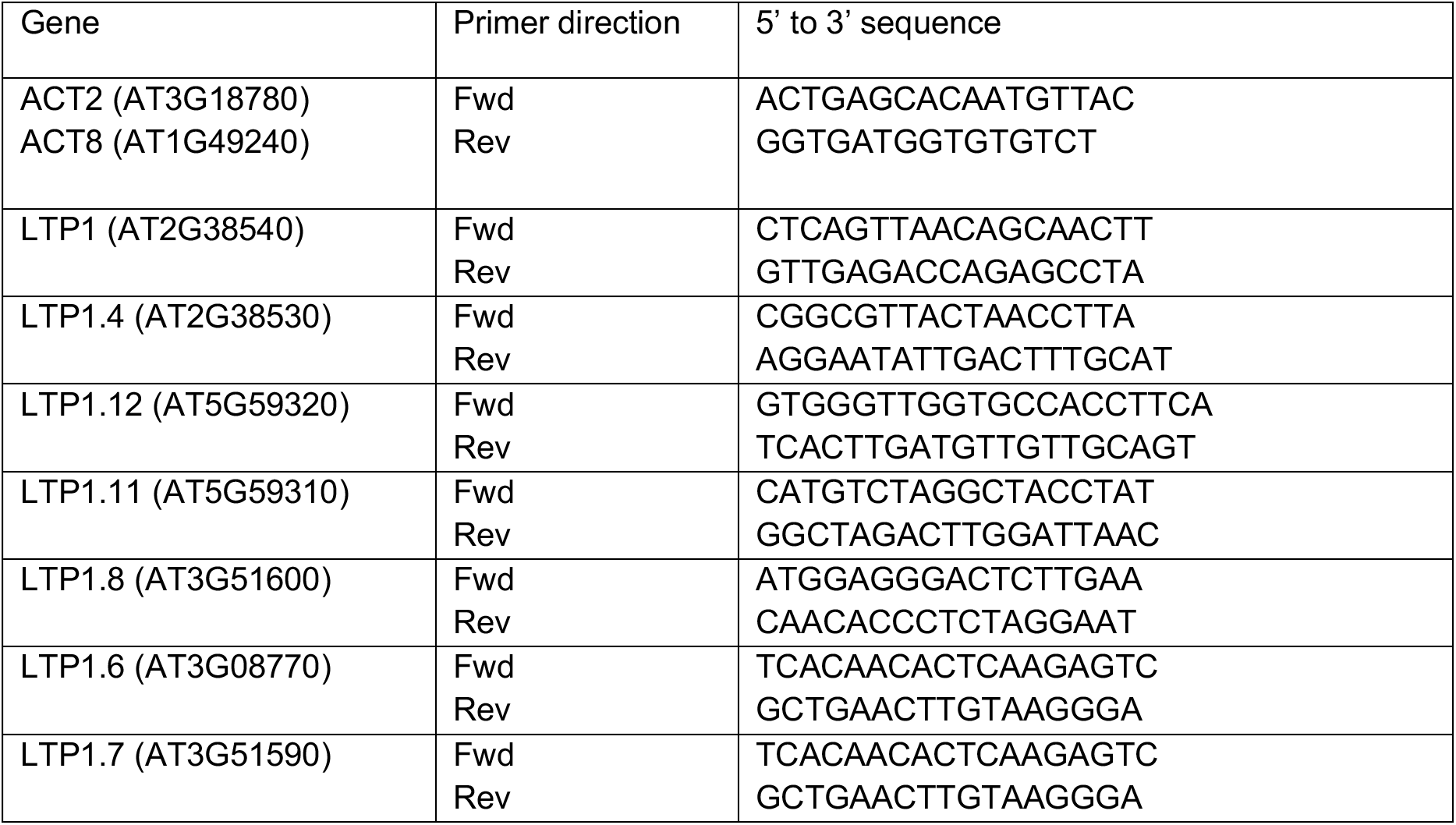
List of qPCR primers.

**Supplemental Table 5.**
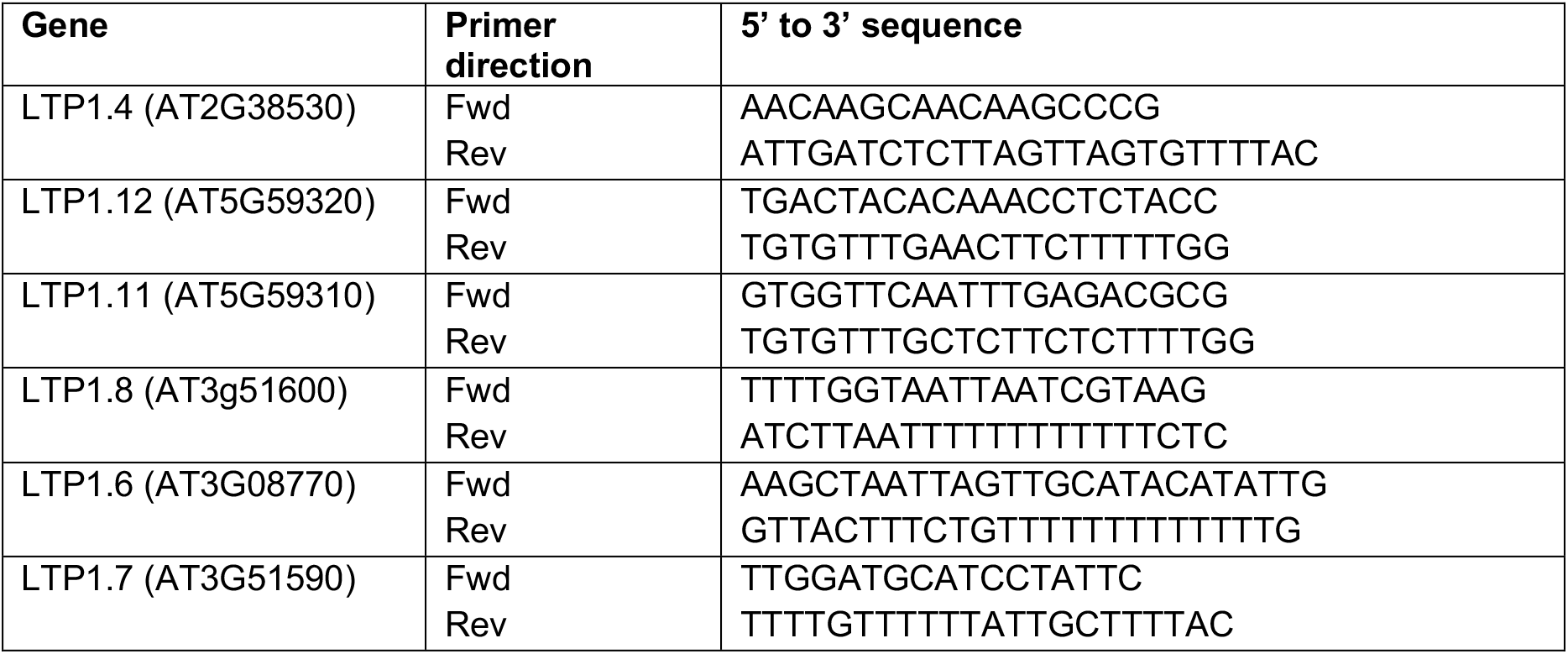
List of primers used for amplifying LTP promoter.

**Supplemental Table 6.**
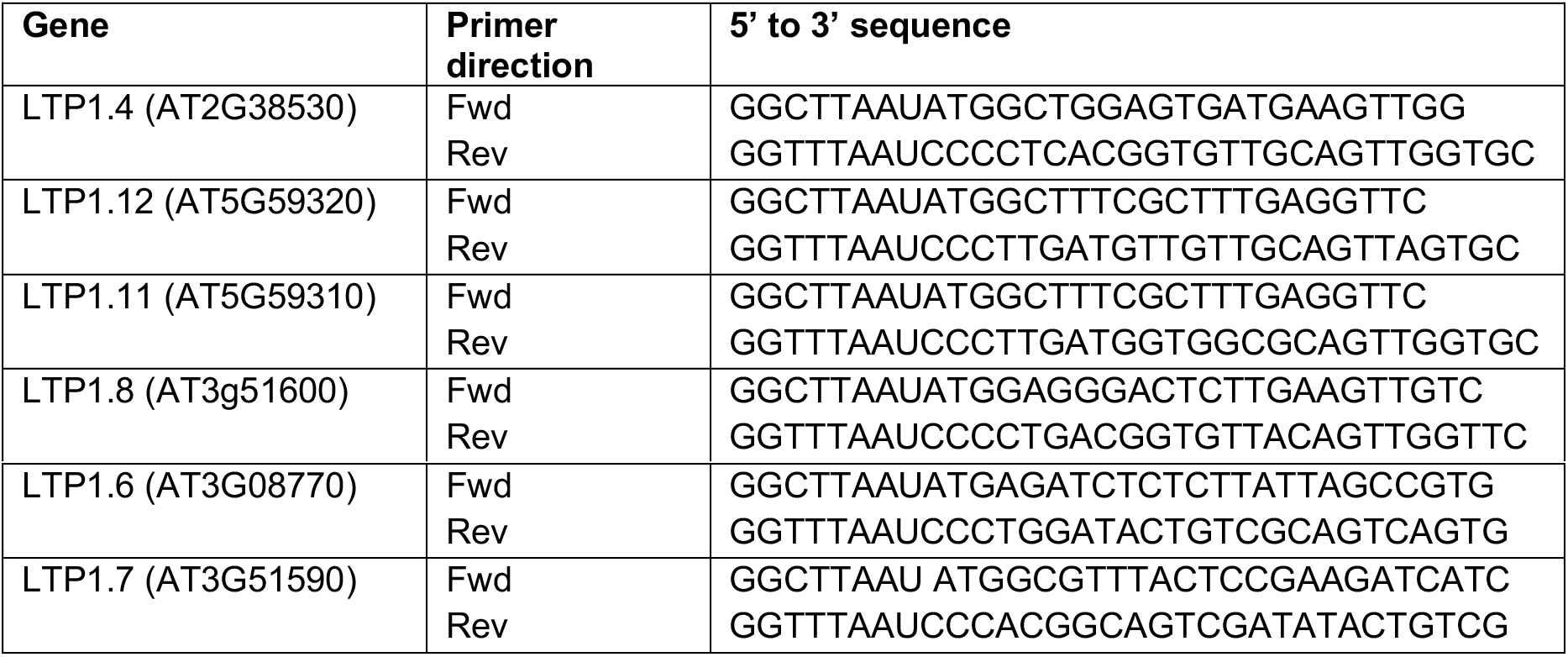
List of primers used for amplifying LTP cDNA.

**Supplemental Table 7.**
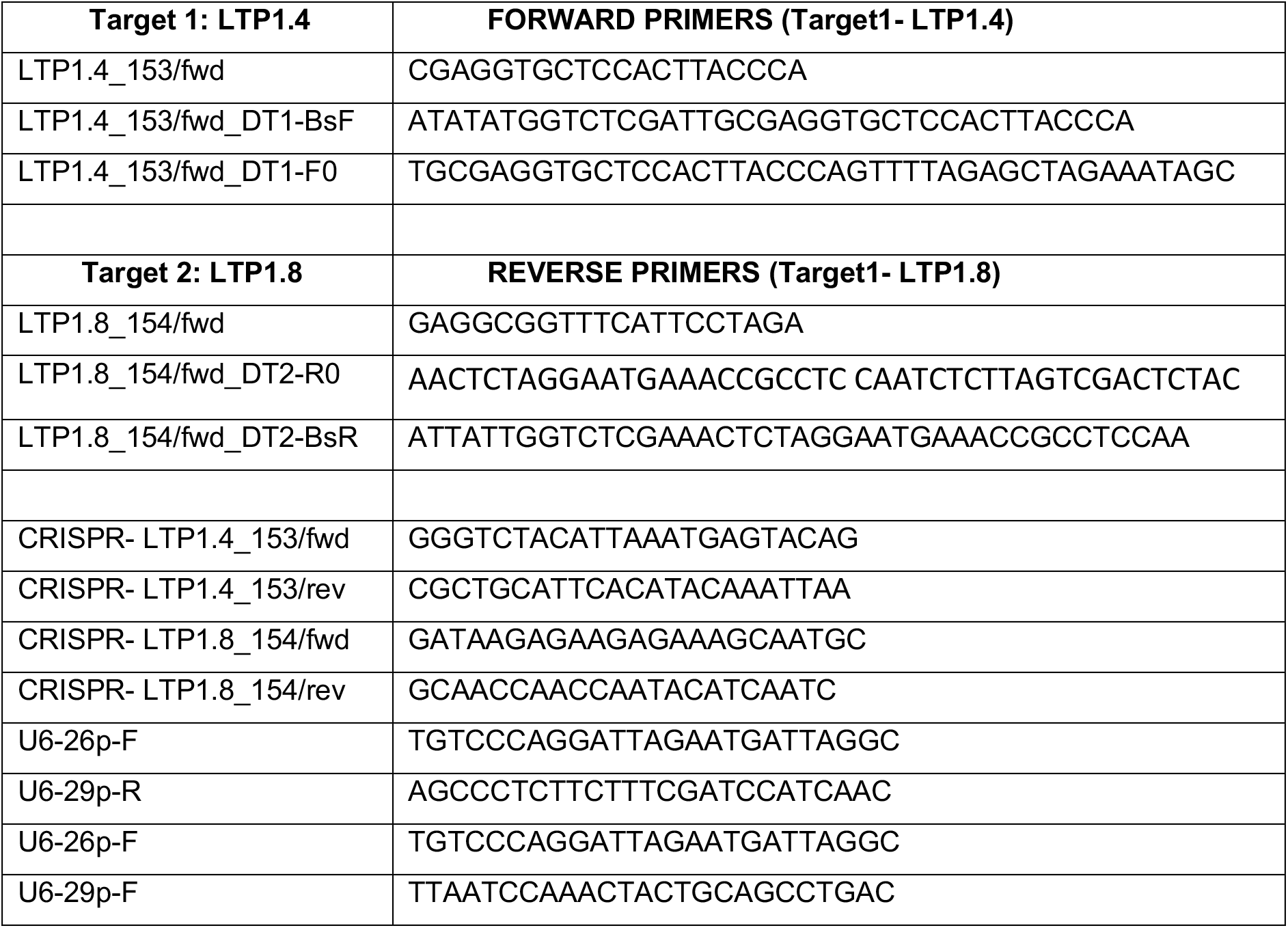
List of primers used for generating LTP1.4sgRNA and LTP1.8sgRNA.

